# Cell type-centric interaction networks define spatial architecture of intrahepatic cholangiocarcinoma

**DOI:** 10.64898/2026.06.02.729644

**Authors:** Hsin-Pei Lee, Meng Liu, Wenqi Wu, Ngoc Thao Lam Nguyen, Jittiporn Chaisaingmongkol, Darko Castven, Ethan Levy, Noemi Kedei, Maria O. Hernandez, Manjari Kundu, Marshonna Forgues, Man-Hsin Hung, Anuradha Budhu, Nada Alani, Stephen M. Hewitt, Ross Lake, Eytan Ruppin, Stanley Lipkowitz, Xin Wei Wang, Jens U. Marquardt, Mathuros Ruchirawat, Lichun Ma

**Author notes:** Correspondence: L.M., M.R., or J.U.M. These authors contributed equally (Hsin-Pei Lee, Meng Liu, Wenqi Wu). <u>Main Contact:</u> Lichun Ma, PhD, Cancer Data Science Laboratory, Center for Cancer Research, National Cancer Institute, NIH, 41 Medlars Drive, Bldg. 41 Rm. A100B, Bethesda, MD 20892.

## Abstract

Tumor spatial organization critically shapes disease progression and therapeutic response, yet remains poorly defined. Intrahepatic cholangiocarcinoma (iCCA), a rare and aggressive liver malignancy with extensive stromal and immune remodeling, provides a compelling model to study tumor architecture. We generated a single-cell spatial atlas of 1 million cells from 131 iCCA patients using 53-plex spatial proteomics. To systemically characterize tumor spatial organization, we developed a graph-based deep learning framework to define cell type-centric interaction networks, identifying 41 distinct multicellular spatial patterns. Integration of these networks revealed higher-order tumor- and immune-enriched microenvironments associated with patient outcomes. Notably, neutrophil-associated tumor-enriched and tumor-desert microenvironments delineated patient groups with opposing clinical outcomes and distinct neutrophil states. These findings were validated by single-cell spatial transcriptomic profiling of 6 million cells from 162 iCCA patients. Together, this study defines the spatial architecture of iCCA and provides a comprehensive resource for exploring tumor spatial organization.

## Introduction

A malignant tumor is a spatially organized ecosystem of diverse tumor, stromal, and immune cell populations, whose interactions critically shape therapeutic response, disease progression, and patient outcome^1,2^. However, resolving this spatial architecture remains a major challenge, as cellular organization within the tumor microenvironment (TME) is highly complex and context-dependent, and cannot be fully captured by conventional profiling approaches^3^. Intrahepatic cholangiocarcinoma (iCCA) provides a compelling model to address this challenge, owing to its highly desmoplastic stroma, extensive immune infiltration, and pronounced cellular heterogeneity, all of which indicate complex spatial organization^4,5^. Clinically, iCCA is a highly aggressive malignancy arising from the intrahepatic bile ducts and represents a rare histological subtype of primary liver cancer. Its global incidence has been steadily increasing, yet the 5-year overall survival remains below 10%, underscoring an urgent need to better understand its tumor biology and improve therapeutic strategies^6,7^.

Efforts in bulk molecular profiling have identified distinct molecular subtypes of iCCA associated with patient outcomes, and single-cell analyses have further refined our understanding by revealing substantial intra- and inter-tumor heterogeneity^5,8–13^. However, these approaches largely lack spatial resolution, and the spatial context in which tumor, immune, and stromal cells interact particularly how they organize into multicellular structures and functional niches, remains poorly defined. Recent advances in imaging-based spatial profiling technologies, including co-detection by indexing (CODEX) and the CosMx spatial molecular imager, have enabled high-dimensional, spatially resolved profiling of tissues at single-cell resolution^14,15^. These approaches have begun to uncover spatially organized tumor and immune niches across multiple cancer types^16–18^, yet systematically defining biologically meaningful spatial structures from large-scale single-cell spatial data remains a fundamental challenge.

Here, we systematically characterize the single-cell spatial landscape of iCCA and identify spatial features associated with patient prognosis. Using CODEX profiling of paired tumor and non-tumor tissues from 131 iCCA patients, we generated a high-dimensional dataset consisting of approximately 1.1 million cells across diverse tumor, stromal, immune, and liver-specific cell types. We then developed a graph-based deep learning framework to define cell type-centric local interaction networks and higher-order spatial organization, uncovering tumor-and immune-enriched microenvironments associated with clinical outcomes. Notably, neutrophil-associated spatial microenvironments delineated patient groups with opposing prognosis and distinct functional states. These findings were independently validated using single-cell spatial transcriptomic profiling of approximately 6 million cells from 162 iCCA patients. Finally, we projected these spatial features into a bulk transcriptome-based scoring framework that stratified patients across independent cohorts and correlated with immunotherapy response. Together, this study establishes a single cell spatial atlas of iCCA and identifies spatially organized microenvironments as key determinants of tumor aggressiveness.

## Results

### Single-cell spatial characterization of iCCA

To characterize the spatial landscape of iCCA, we performed single-cell spatial proteomic profiling of the Thailand Initiative in Genomics and Expression Research for Liver Cancer (TIGER-LC) cohort as the discovery cohort, and single-cell spatial transcriptomic profiling of an independent European cohort for validation (Figure 1A). In the TIGER-LC cohort, paired tumor and adjacent non-tumor tissues were collected from 131 Thai patients undergoing surgical resection (Table S1). Most patients were over 50 years old (85%) and negative for hepatitis B (79%) and hepatitis C (82%). We constructed separate tissue microarrays (TMAs) of tumor and non-tumor tissues and performed CODEX-based spatial proteomics at single-cell resolution using a curated 53-plex antibody panel (Tables S2 and S3). For the European validation cohort, we collected 42 tumor samples from 31 iCCA patients (two samples from 11 patients and one sample from 20 patients, Table S4), including 6 samples from 3 previously reported cases^19^. Single-cell spatial transcriptomic profiling was performed using the CosMx platform. Unlike the TMA-based discovery cohort, the CosMx data were generated from whole-slide tissue sections, enabling broader spatial coverage and reducing sampling bias. In total, 21 CosMx slides were imaged. This dataset served as a cross-cohort and cross-modality resource to validate the findings from the TIGER-LC discovery cohort.

**Figure 1.**
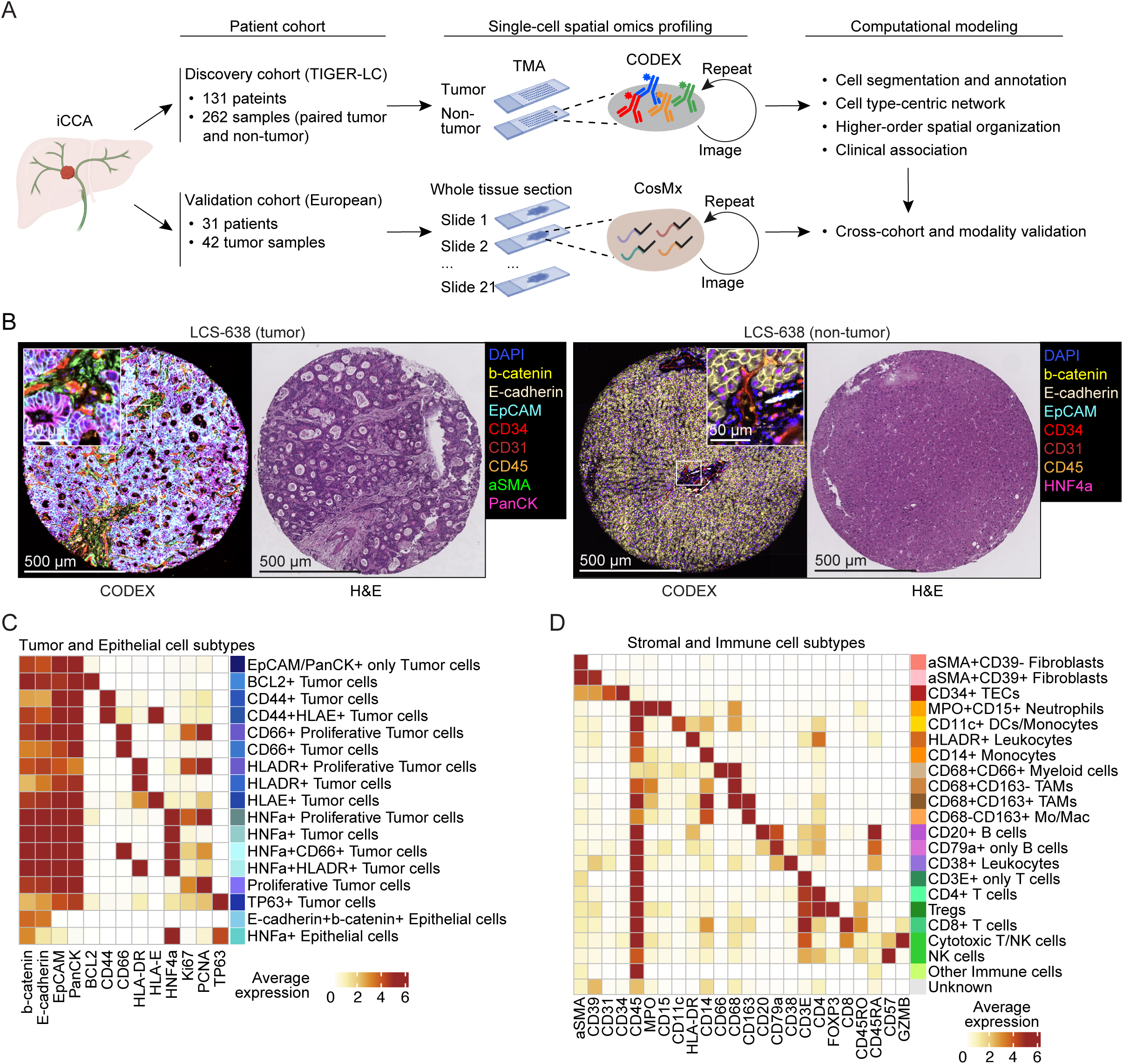
Characterization of the iCCA single-cell spatial landscape. (A) Overview of the workflow. Tumor and paired non-tumor samples from 131 iCCA patients in the TIGER-LC cohort (discovery cohort) were profiled using CODEX with TMA. For the validation cohort, 42 tumor samples from 31 iCCA patients in the European cohort were subjected to whole-tissue slide single-cell spatial transcriptomic profiling using the CosMx platform (38 samples with 1K gene panel and 4 samples with 6K gene panel). In total, 21 CosMx slides were profiled, with two samples per slide. (B) Representative tumor (left) and non-tumor (right) shown by CODEX image and H&E for one patient (LCS-638). (C and D) Heatmap of the average marker expression for tumor and epithelial cell subtypes (C) and stromal and immune cell subtypes (D) in tumor samples. TEC, tumor-associated endothelial cell; TAM, tumor-associated macrophages; DC, dendritic cell; NK, natural killer cell; Mo/Mac, monocyte-like or macrophage-like.

The CODEX marker panel enabled characterization of tumor cells, hepatocytes, biliary epithelial cells, various immune populations, stromal components, and functional states such as proliferation (Table S3). The CODEX staining demonstrates high quality and corresponds well with histological features observed in H&E images. For example, CODEX captures tumor structures (EpCAM) and vasculatures (CD31/CD34) surrounded by fibroblasts (αSMA) in tumor tissues (Figure 1B). In non-tumor samples, CODEX reveals plates of hepatocytes (HNF4α), and portal triad with artery (CD31/CD34) and bile duct (b-catenin/E-cadherin/EpCAM) (Figure 1B). After cell segmentation and quality control (Methods), we annotated cell types and subtypes of 544,131 cells from tumor samples using known marker expression profiles (Figures 1C, 1D, S1A, and S1B; see Methods for details). Within the epithelial population (b-catenin+E-cadherin+CD45-) in tumor samples, cells expressing EpCAM and/or PanCK were classified as tumor cells, while those lacking both markers were classified as epithelial cells. Further subtype annotation using additional markers identified 17 tumor and epithelial subtypes, including proliferative (Ki67+/PCNA+) tumor cells, HLADR+ tumor cells, among others (Figure 1C). In the TME, we identified 2 subtypes of cancer-associated fibroblasts (CAFs, αSMA+CD39+/-), tumor-associated endothelial cells (TECs, CD31+/CD34+), 8 subtypes of myeloid cells and 9 subtypes of lymphocytes (Figure 1D). Spatial mapping of annotated cell types confirmed concordance with CODEX staining (Figures S1C). Similarly, we analyzed 577,473 cells from non-tumor samples after quality control and identified 22 cell subtypes (Figures S1D-S1F). Fifteen cell subtype annotations, primarily immune and stromal cells, were shared between tumor and non-tumor samples, while others were unique to either tumor or non-tumor tissues.

Cell type composition analysis showed that non-tumor tissues were predominantly composed of hepatocytes (Figure S1G). In contrast, tumor samples were significantly enriched for EpCAM+/PanCK+ tumor cells compared with a small proportion of biliary epithelial cells in non-tumor (*p*<0.0001; Figure S1G). Tumor samples also showed marked expansion of stromal, lymphoid and myeloid compartments. Upon closer examination of the immune cells, we found substantial expansion of the MPO+CD15+ neutrophil population in tumor tissues, alongside increases in CD68+CD163- macrophages, CD4+ T cells, CD38+ leukocytes, CD8+ T cells and CD20+ B cells (Figure S1H). Conversely, reduced levels of CD68+CD163+ macrophages and HLADR+ leukocytes were observed in tumor samples (Figure S1H).

Together, these results establish a high-resolution single-cell spatial atlas of iCCA and highlight pronounced cellular heterogeneity.

### Mapping multicellular interaction networks to delineate the spatial organization of iCCA

A systemic characterization of the multicellular spatial interaction networks may advance our understanding of cellular organization in iCCA. Here, we followed a framework where tissue architecture is built from elementary units of local cellular interaction networks, which organize into higher-order cellular structures to support complex biological functions. Accordingly, we employed a two-level strategy to delineate the spatial architecture of iCCA: first, we defined local cellular organization by identifying cell type-centric multicellular networks representing the immediate cellular neighborhoods of individual cell types as elementary building blocks of higher- order structure; second, we resolved higher-order spatial organization by integrating these local networks (Figure 2A). We captured these local networks using spatial dynamics networks (SDNs), which systematically characterize the composition and interactions within the local microenvironment surrounding each cell type (Figure 2A). Building on our previous SDN framework for tumor cell states^19^, we extend this concept to all cell types to enable a comprehensive reconstruction of tumor spatial organization. We applied graph attention networks (GATs) to capture the intricate cellular interactions based on the spatial proximity of cells, by leveraging both cell (sub)type information (subtype information of tumor cells was omitted to reduce model complexity) and neighborhood graphs of individual cells constructed from their spatial coordinates. A total of 41 SDNs were derived, including 8 tumor cell SDNs (SDN_T), 9 TEC SDNs (SDN_E), 6 CAF SDNs (SDN_F), 10 myeloid cell SDNs (SDN_M), and 8 lymphocyte SDNs (SDN_L). We further resolved higher-order cellular organizations by learning SDN spatial co-localization patterns using the GAT method (Figure 2A). This analysis revealed seven higher-order cellular organizations termed super-SDNs (sSDNs) and each comprising a distinct combination of SDNs. The two-level strategy enabled multiscale characterization of tumor spatial organization, providing a comprehensive view of global tumor architecture.

**Figure 2.**
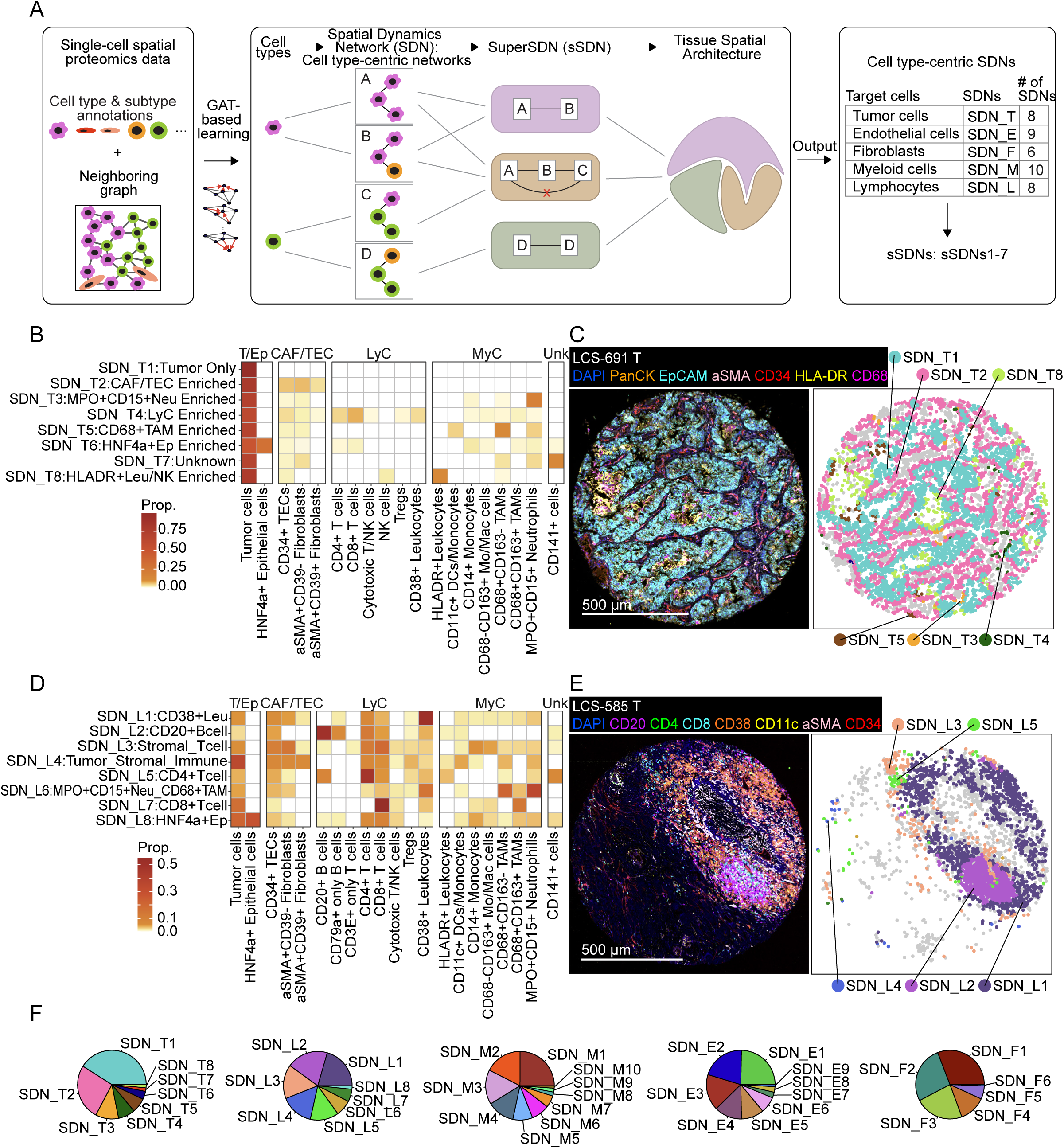
Mapping multicellular spatial organization by defining SDNs and sSDNs. (A) Schematic overview of SDN and sSDN characterization. Cell (sub)type information and neighborhood graphs of individual cells constructed from their spatial coordinates were used as input for the GAT. The learned embeddings from GAT were used for clustering to identify SDNs in a cell-type-centric manner and further named based on neighbor cell type composition. Higher-order cellular organizations were determined by integrating neighboring SDNs using the GAT-based approach. The summary table displays the 41 identified SDNs and 7 sSDNs. (B) Heatmap of SDN_T cell subtype composition. All tumor subtypes were combined as “Tumor cells.” Cell subtypes with greater than 0.01 proportion are shown. (C) A representative tumor sample showing tumor-only area and tumor-stromal interface in CODEX staining (left) and SDN annotation (right). (D) Heatmap of SDN_L cell subtype composition. Cell subtypes with greater than 0.01 proportion are shown. (E) A representative tumor sample with a TLS-like structure in CODEX staining (left) and SDN (right) annotation. (F) Pie plots of the SDN proportions within each cell type category. (B and D) T/Ep, tumor cell/epithelial cell; CAF/TEC, cancer-associated fibroblast/tumor-associated endothelial cells; LyC, lymphocyte; MyC, myeloid cell; Neu, neutrophil; Leu, leukocyte; Unk, unknown.

In the assessment of tumor cells, SDNs effectively capture diverse tumor cell neighborhoods (Figures 2B and S2A). Among these tumor cell SDNs (SDN_T), we identified distinct network types, including a bulk tumor network (SDN_T1) and a tumor-stromal network (SDN_T2) and four tumor-immune networks characterized by enrichment of tumor cells with MPO+CD15+ neutrophils (SDN_T3), CD8+ and CD4+ T cells (SDN_T4), CD68+CD163-TAMs (SDN_T5), and HLADR+ leukocytes (SDN_T8). In addition, SDN_T6 was enriched for epithelial cells lacking tumor markers, while SDN_T7 showed no clear enrichment for specific lineage-defined cell types. Across all SDN_Ts, tumor cells remained the predominant component (Figure 2B). Mapping SDN_T assignments to spatial coordinates revealed distinct localization patterns. For example, SDN_T1 frequently interacted with SDN_T2, delineating tumor-stromal boundary regions (Figure 2C). Tumor cell subtypes also exhibited distinct spatial preferences, as demonstrated by enrichment analysis across SDN_Ts (Figure S2B). Notably, HLADR+ tumor cells were preferentially localized to immune-enriched networks such as SDN_T8 and SDN_T4 (Figures S2B and S2C). These results suggest that SDN_Ts successfully capture multicellular spatial networks centered on individual tumor cells, revealing diverse interaction patterns between tumor cells and the TME.

The SDNs also captured immune and stromal tissue structures. Eight lymphocyte SDNs (SDN_L) were identified (Figures 2D and S2D). Compared with tumor SDNs, these showed more diverse neighboring cell type compositions, although several were characterized by a high abundance of specific cell types, such as SDN_L1 (CD38+ leukocytes), SDN_L2 (CD20+ B cells), SDN_L5 (CD4+ T cells), and SDN_L7 (CD8+ T cells) (Figure 2D). Notably, SDN_L2 showed strong co-localization of CD20+ B cells with CD4+ T cells and CD8+ T cells, and mapped to a spatial structure resembling a tertiary lymphoid structure (TLS; Figure 2E). This result suggested that the SDN-based approach could capture known multicellular organizations with meaningful biological functions. Similarly, we obtained nine TEC SDNs (SDN_E) and six CAF SDNs (SDN_F; Figures S2E-S2G). Finally, we identified ten myeloid cell SDNs (SDN_M; Figures S2H and S2I). Noticeably, MPO+CD15+ neutrophils were strongly enriched in three SDN_Ms, each highlighting distinct spatial patterns: CD68+ TAM and MPO+CD15+ neutrophil co-localization in SDN_M1, MPO+CD15+ neutrophil aggregation in SDN_M2, and tumor–MPO+CD15+ neutrophil interactions in SDN_M4. Specifically, SDN_M1 had uniquely high proportions of CD38+ leukocytes, while SDN_M4 contained immunosuppressive CD68-CD163+ Mo/Mac cells (monocyte-like or macrophage-like cells) (Figure S2H). CD38 expression has been reported in tumor-infiltrating lymphocytes and can promote inflammatory anti-tumor response^20^. It has also been reported to be expressed by M1 macrophages and neutrophils associated with T cell-mediated anti-tumor response^21^. Thus, SDN_M1 may represent inflamed TME with active anti-tumor immune response. By contrast, SDN_M4 may represent an immunosuppressive environment. Collectively, a total of 41 SDNs were identified across 5 main cell types in iCCA tumors (Figures 2F and S2J). These SDNs could efficiently capture both known and novel biologically meaningful multicellular interaction networks, comprehensively defining the landscape of cell type-centric local organizations in iCCA.

### Higher-order cellular organizational structures are linked to patient outcomes

To understand higher-order cellular organizations, we integrated neighboring SDNs using a GAT-based approach (Figure 2A; Methods). Unsupervised clustering of the resulting embeddings yielded ten clusters, three of which were removed from further analysis due to low cell numbers or enrichment for epithelial cells lacking tumor markers (Figures S3A-S3C; Methods). Seven distinct higher-order multicellular organizations, i.e., sSDNs, were derived (Figure 3A). we found that five sSDNs were predominantly tumor-associated (Figures 3A and S3C). Specifically, sSDN5 consisted almost entirely of tumor cells, characterizing regions with uniform tumor cell co-localization. Other tumor-enriched sSDNs captured interfaces between tumor and non-tumor cells. The sSDN1 (Tumor_Stromal) was enriched for SDN_T1 and SDN_T2, indicating tumor-stromal co-localization. Interaction analysis showed that SDN_T1 interacted with SDN_T2, and SDN_T2 interacted with SDN_F1, whereas SDN_T1 did not directly interact with SDN_F1, suggesting a spatial relationship of bulk tumor region, tumor-stromal interface, and stromal cells. The sSDN2 (Tumor_Myeloid) is characterized by enriched interactions between SDN_T3 and SDN_M4, indicating tumor-neutrophil crosstalk. The additional high proportions of SDN_T1, SDN_T2, SDN_T5 further suggested the infiltration of neutrophils and TAMs in these tumor regions. The sSDN6 (Tumor_Stromal_Myeloid) is centered around SDN_T7 interaction with other SDNs such as SDN_M5 and SDN_T1. Lastly, sSDN7 (Tumor_HLADR+Leu_NK) showed strong co-localization of SDN_T8 and SDN_M6, with SDN_T8 mediating interactions with other tumor SDNs, representing tumor cell regions with HLADR+ leukocyte and NK cell infiltration. Among the tumor-enriched sSDNs, sSDN1 was the most abundant across samples, suggesting that our approach captured the glandular and fibrous stroma iCCA phenotype^22^ (Figure S3B). Notably, not all SDNs interacted within each sSDNs, and individual SDNs may contribute to multiple sSDNs, highlighting the importance of spatial context in defining the iCCA landscape.

**Figure 3.**
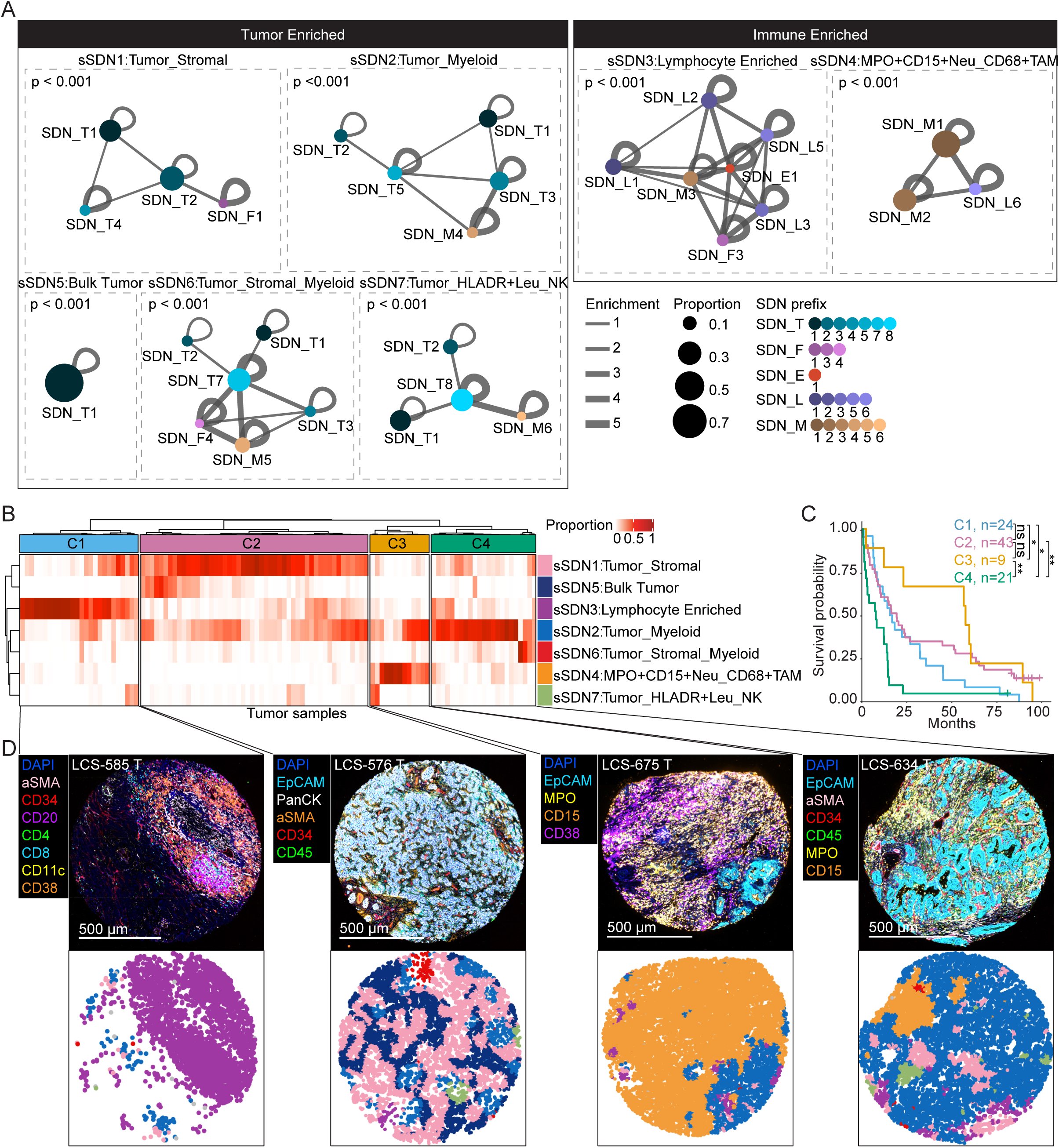
sSDNs are associated with patient outcomes. (A) The sSDNs represented by SDN-SDN co-localizations. Only SDNs with greater than 0.05 proportions in a sSDN are shown. Node size indicates proportion. Line thickness indicates co-localization enrichment. The nodes are colored by SDN. Statistical significance of each sSDN structure was calculated using the Hotelling’s *T*-squared test between SDN composition of each structure compared with the averaged SDN composition in each sample. For sSDN5 composed of one SDN, *p*-value was calculated with two-sided Student’s *t*-test. (B) Hierarchical clustering of tumor samples by sSDN compositions in each sample using Pearson correlation distance and “ward.D2” method. (C) Overall survival of the four iCCA patient groups from (B). Pairwise *p-*value was calculated using log rank test. \**p<*0.05, \*\**p*<0.01. (D) Representative CODEX images (top) and sSDNs (bottom) mapped to spatial coordinates of the four patient groups. See (B) for sSDN color coding.

In addition to tumor-enriched sSDNs, we identified two immune aggregates with minimal tumor content (Figures 3A and S3C). The sSDN3 (Lymphocyte Enriched) included a mixture of various lymphocytic SDNs, including SDN_L1, SDN_L2, and SDN_L3. By contrast, sSDN4 (MPO+CD15+Neu_CD68+TAM) showed high proportions of myeloid SDNs enriched for MPO+CD15+ neutrophils, CD68+ TAMs, and CD38+ leukocytes. As described earlier, the association of these cells may suggest an inflammatory immune environment. Spatial analysis revealed strong spatial aggregation of sSDNs, as indicated by Moran’s I values. Individual SDNs showed varying degrees of spatial autocorrelation, with endothelial-related SDNs being relatively dispersed and tumor-associated SDNs more clustered (Figures S3D and S3E). Collectively, defining sSDNs allowed us to capture complex tissue structures involving tumor cells, stromal cells, and immune cells.

To understand the biological implications of the sSDNs, we stratified iCCA patients into four groups based on hierarchical clustering of sSDN proportions in each sample and analyzed the overall patient survival (Figures 3B and 3C). Patient cluster C1 was dominated by sSDN3 with low tumor cell presence and high co-localizations among T cells, CD38+ leukocytes, CD20+ B cells, and stromal cells (Figures 3B and S3C). C2 was dominated by sSDN1, indicating tumors with high stromal components and little immune presence. C3 had uniquely high proportion of sSDN4, suggesting innate immune activity (Figures 3B and S3F). Lastly, C4 had high proportions of sSDN2 and sSDN6, indicating tumor-myeloid interactions (Figures 3B and S3G). These four patient clusters showed strong prognosis stratification (overall *p*=0.00084), with C3 related to the most favorable survival, C4 linked to the poorest outcomes, and C1 and C2 in between (Figure 3C). Representative tumor samples in each patient group showed concordance between sSDN definition and CODEX staining of cells (Figures 3D and S3H). The survival associations remained consistent when restricting analyses to patients with similar tumor cellularity (the number of tumor cells per unit area; Figures S4A-S4C). In contrast, the overall survival of these iCCA patients could not be significantly stratified based on cell (sub)type composition of the tumor samples or simply tumor-immune ratio, underscoring a need to study iCCA TME in the spatial context (Figures S4D-S4F). We did not observe significant associations between patient clusters with known iCCA driver mutations or clinical variables in our cohort (Figures S4G and S4H).

Collectively, by learning higher-order spatial organizations, we identified complex multicellular organizations related to patient prognosis. Specifically, C3 and C4, representing the best and worst patient prognosis groups, were related to myeloid cell spatial organizations, underscoring the need to further understand myeloid cells based on their spatial locations.

### Validation of spatial cellular organization in an independent cohort by single-cell spatial transcriptomics

Because the TIGER-LC cohort was profiled using TMA-based spatial proteomics with limited coverage, we sought to assess whether SDN and sSDN frameworks generalize to whole-tissue sections profiled at higher molecular resolution. To this end, we performed single-cell spatial transcriptomic profiling of iCCA in an independent European cohort using the CosMx platform (Figure 1A; Table S4). After quality control, we obtained 5,558,983 high-quality cells across 42 tumor samples from 31 patients, spanning immune, stromal, malignant, and non-malignant epithelial populations, with detailed subtype annotations (Figures 4A, 4B, and S5A-S5E). Transcriptome-based cell type annotations were highly concordant with protein staining and histological features, supporting the quality and reliability of the dataset (Figure 4C). To infer spatial organization, we trained an XGBoost model on CODEX-data from the TIGER-LC cohort and applied it to predict SDNs and sSDNs in the European cohort (Methods). Most SDNs (90.2%) and all sSDNs were reproducibly detected across multiple samples (≥3) in the European cohort, demonstrating robust cross-cohort and cross-modality generalizability (Figures 4D and 4E). The SDN composition within each predicted sSDN closely recapitulated patterns observed in the TIGER-LC cohort (Figure S5F).

**Figure 4.**
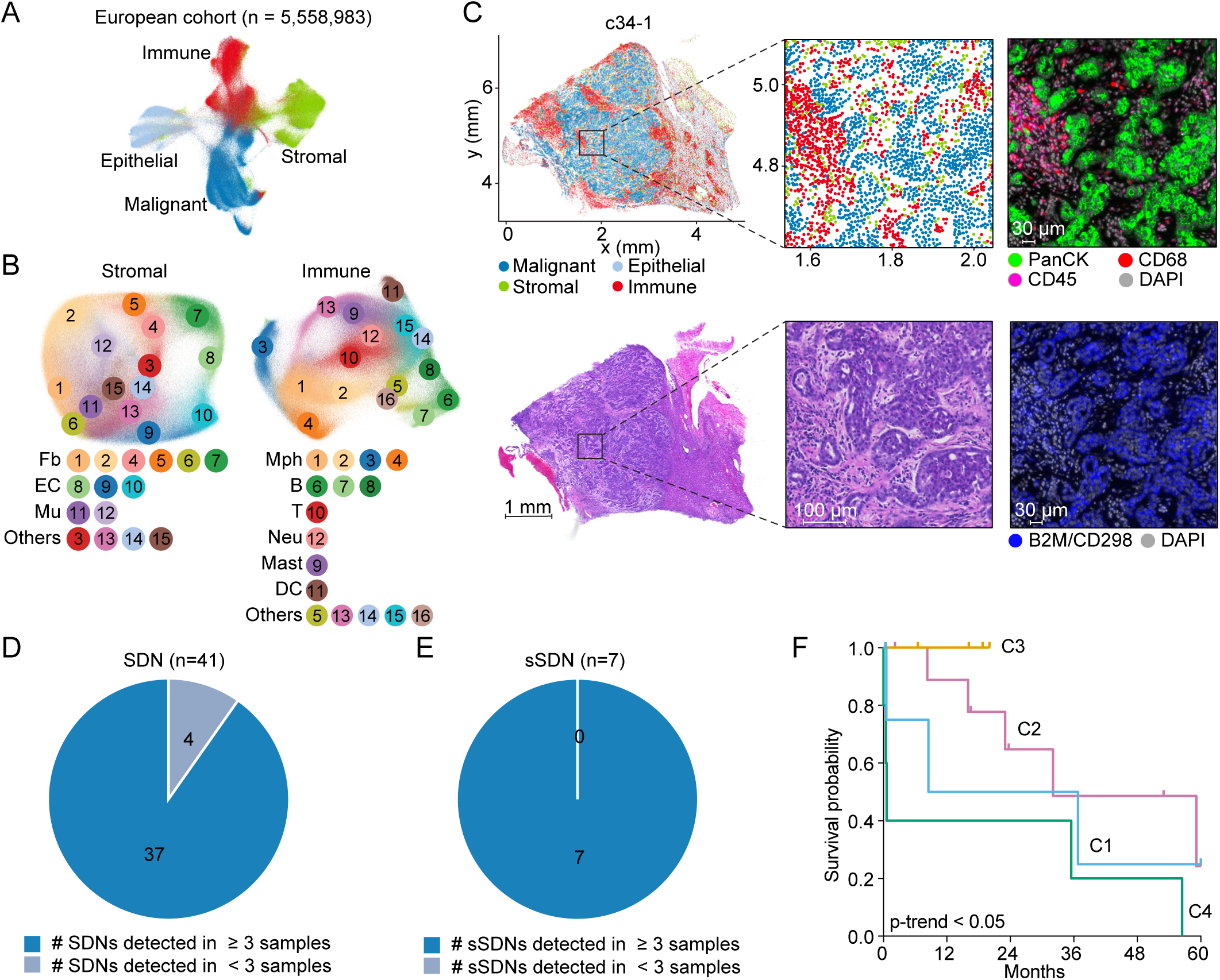
Single-cell spatial transcriptome validation of spatial cellular organizations in a European iCCA patient cohort. (A) UMAP of 5,558,983 single cells after quality control, with different cell types indicated by colors. (B) UMAP of stromal (left) and immune (right) cell clusters. Detailed annotations were shown in Figure S5. Fb, fibroblast; EC, endothelial cell; Mu, mural cell; Mph, macrophage; B, B cell; T, T cell; Neu, neutrophil; DC, dendritic cell. (C) Cell type annotation, protein staining and histology image for a representative sample (c34-1). CD68 (red), CD45 (magenta) and Pan-cytokeratin (Pan-CK, green) represent markers for macrophages, immune cells, and epithelial cells, respectively. DAPI (light grey) and B2M/CD298 (blue) were used for nuclei and membrane staining. (D and E) The number of SDNs (D) and sSDNs (E) detected in the European cohort. (F) Kaplan-Meier plot of the five-year survival of four iCCA patient groups in the European iCCA cohort. For patients with two samples collected, the patient group was assigned based on the highest correlation value. *p*-trend was calculated using log rank test for trend.

We further assigned each sample to patient clusters based on similarity in sSDN enrichment to the four patient clusters (C1-C4) identified in the TIGER-LC cohort. Survival analysis revealed consistent stratification, with C3- and C4-like groups showing the best and worst outcomes, respectively, and C1- and C2-like groups displaying intermediate survival (Figure 4F). Together, these results validate the SDN- and sSDN-defined spatial architectures and their clinical relevance across independent cohorts and spatial data modalities.

### MPO+CD15+ neutrophil-related environments underly survival differences in iCCA

To elucidate the factors underlying the distinct survival outcomes of C3 and C4 patient clusters, we analyzed their predominant sSDNs in the TIGER-LC cohort. The proportion of sSDN2 was significantly higher in C4 (*p<*0.001; Figures 3B and 5A), while sSDN4 was significantly enriched in C3 (*p*<0.0001; Figures 3B and 5B). Compositional analysis revealed that sSDN2 consisted mainly of SDN_T (66%) and SDN_M (21%), whereas sSDN4 was dominated by SDN_M (80%) with minimal SDN_T (4.5%) involvement (Figure 5C). Direct quantification of tumor and immune cell proportions confirmed a higher tumor cell fraction in C4 and a higher immune cell fraction in C3 (Figures S6A and S6B). To further characterize the differences at the molecular level, we analyzed bulk transcriptome data of 91 patients in this cohort generated from our previous study^11^. Differential gene expression analysis revealed that C4 was enriched for genes associated with tumor progression (e.g., *ASPN, SPP1*) and immunosuppression (e.g., *CD276*), whereas C3 showed upregulation of genes involved in innate immunity and inflammation (e.g., *CLEC4E, CLEC4D, CXCR2, CXCL2, CSF3R, TNF*; Figure S6C). Notably, *CSF3R* plays a crucial role in neutrophil proliferation and differentiation^23^, and *CXCR2* is a chemoattractant that mediates neutrophil release from bone marrow and migration to inflamed sites^24^. Pathway enrichment analysis revealed upregulation of pro-tumor pathways in C4, including epithelial-mesenchymal transition, glycolysis, and KRAS signaling, while C3 showed enrichment of inflammatory pathways (Figure S6D). Together, these results suggest that poor prognosis in C4 may stem from increased pro-tumor signaling associated with tumor-myeloid interactions and reduced anti-tumor inflammatory responses.

**Figure 5.**
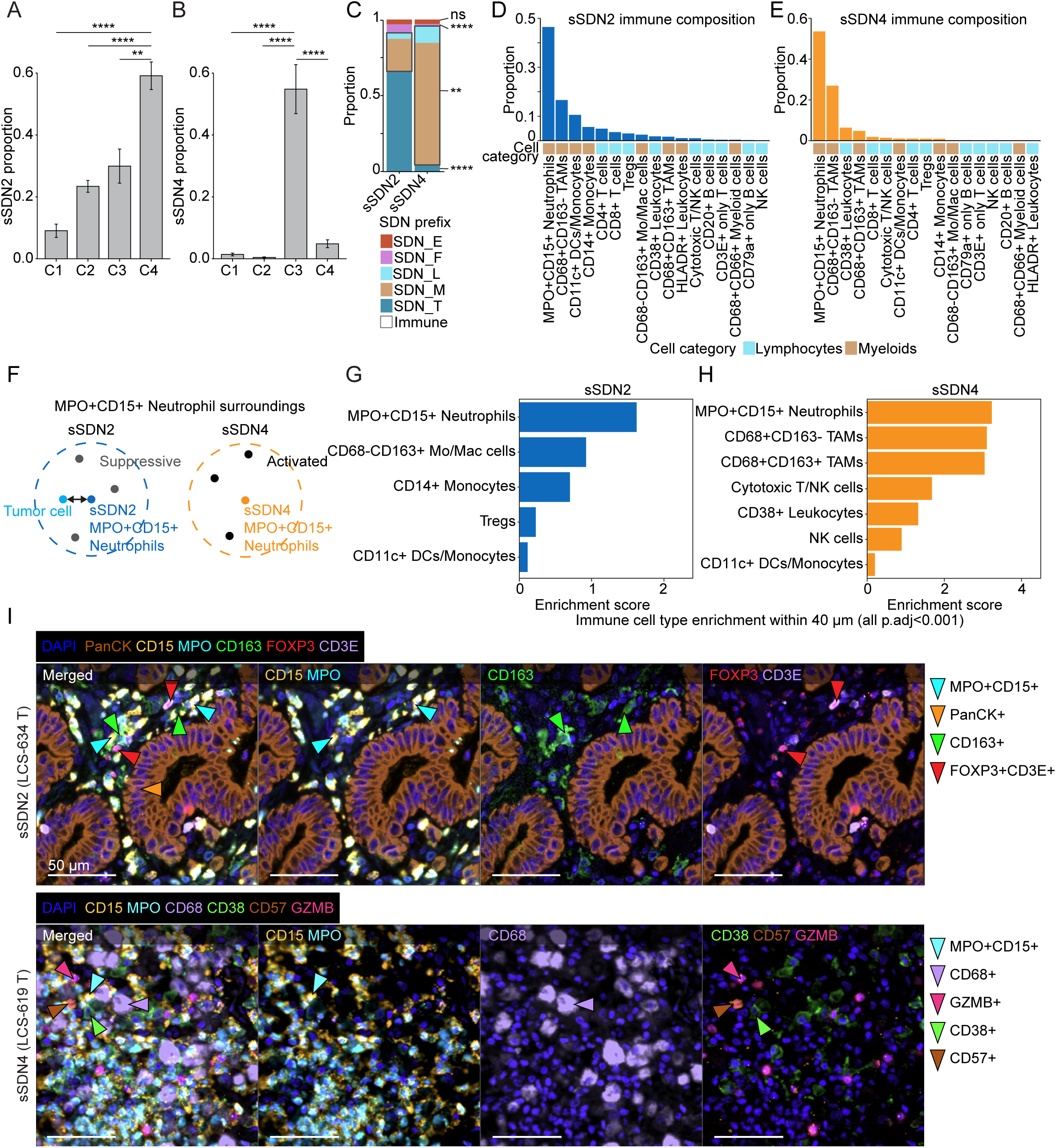
Two distinct MPO+CD15+ neutrophil-associated spatial environments. (A and B) Bar plots of the sSDN2 (A) and sSDN4 (B) by-sample proportions in the four patient clusters. Bar and whiskers indicate the mean and standard error. *p*-values calculated using two-sided Student’s *t*-test in reference to C4 (A) and C3 (B). \*\**p*<0.01, \*\*\*\**p*<0.0001. (C) Stacked bar plot of the overall compositions of SDNs in sSDN2 and sSDN4. The immune cell SDN proportion is outlined in grey box. *p*-values between the same SDN types of the two sSDNs were calculated using two-sided Student’s *t*-test and two immune SDNs were combined for the test. \*\**p*<0.01, \*\*\*\**p*<0.0001, ns, not significant. (D and E) Bar plots of the sSDN2 (D) and sSDN4 (E) immune cell compositions in proportions out of all immune cells in each sSDN. Immune cell types are ranked by decreasing proportion. (F) Illustration of sSDN2 and sSDN4 MPO+CD15+ neutrophil surroundings centered around tumor-interacting sSDN2 neutrophils and sSDN4 neutrophils, respectively. (G-H) The significant immune cell subtype enrichment within 40 µm of sSDN2 MPO+CD15+ neutrophils (G) and sSDN4 MPO+CD15+ neutrophils (H). Enrichment score represents the log2 fold change of the observed immune proportion surrounding each target cell over a randomly permutated proportion, averaged across 1000 permutations. All FDR-adjusted *p*-values < 0.001. (I) Representative CODEX images of sSDN2 MPO+CD15+ neutrophil (top) and sSDN4 MPO+CD15+ neutrophil (bottom) surroundings. Arrows indicate example cells expressing indicated markers.

To gain insight into myeloid cells, we further examined immune cell composition within sSDN2 and sSDN4. Noticeably, MPO+CD15+ neutrophils constituted the largest immune cell fraction in both networks (Figures 5D and 5E). Thus, we hypothesized that the concurrence of MPO+CD15+ neutrophils with tumor cells in sSDN2 may contribute to a neutrophil-mediated immunosuppressive environment, while MPO+CD15+ neutrophils in the absence of tumor cells in sSDN4 may be associated with an immune-activating context (Figure 5F). The divergent MPO+CD15+ neutrophil-related environments may underlie the survival differences between the C4 and C3 clusters. To test the hypothesis, we characterized the enriched immune cell populations surrounding MPO+CD15+ neutrophils based on a 40 µm radius. We found unique enrichments of CD68-CD163+ Mo/Mac cells, CD14+ monocytes and Tregs in the environment of MPO+CD15+ neutrophils within sSDN2 (Figure 5G). CD163+ is a well-known marker for immunosuppressive myeloid cells, and Tregs have established roles in immunosuppression and correlation with poor survival in many cancer types^25–27^. In contrast, CD68+CD163+/- TAMs, cytotoxic T/NK cells, NK cells and CD38+ leukocytes were uniquely enriched in sSDN4 MPO+CD15+ neutrophil surroundings (Figure 5H). Most of these populations have been implicated in anti-tumor immunity and favorable prognosis^28–32^. Notably, pathway enrichment analysis showed the enrichment of TNF-a signaling via nuclear factor kappa-B (NF-kB) in C3 patient cluster (Figure S6D). This pathway is crucial in the polarization of M1 macrophages exhibiting anti-tumor functions^32,33^, aligning with the enrichment of CD68+ TAMs surrounding sSDN4 MPO+CD15+ neutrophils. CODEX images corroborated the spatial co-localization of distinct immune cells with MPO+CD15+ neutrophils associated with the two sSDNs (Figure 5I). Functional protein marker analysis revealed several markers suggesting inflammatory response in the sSDN4 MPO+CD15+ neutrophil surroundings, including elevated CD11b expression in MPO+CD15+ neutrophils, and high IFNG expression by CD38+ leukocytes (Figure S6E). To assess spatial specificity of the enriched immune cells surrounding MPO+CD15+ neutrophils, we quantified the densities of these cells at increasing radii to MPO+CD15+ neutrophils (Figure S6F). Most immune cell densities sharply declined with increased distance, indicating that the cell-cell interactions likely depend on close physical proximity (Figures S6G and S6H).

To investigate molecular signaling in MPO+CD15+ neutrophils associated with sSDN2 and sSDN4, we leveraged scRNA-seq data, which provides higher transcriptomic resolution than spatial platforms. Specifically, we integrated CODEX data with scRNA-seq data from a published liver cancer study^10^ using MaxFuse^34^ (Methods) to identify neutrophils corresponding to sSDN2 and sSDN4. Analysis of neutrophil-related genes indicated that neutrophils in sSDN2 displayed elevated expression of genes such as *DUSP1* and *CD55* (Figure S7A). *DUSP1* has been well linked to anti-inflammation and tumor progression^35,36^. Additionally, *CD55* has been recently shown as a key regulator used by neutrophils in resolving inflammation^37^. By contrast, neutrophils in sSDN4 exhibited elevated expression of genes including *CD83* and *CCL3* which were associated with activated neutrophils and antitumor responses^38,39^ (Figure S7A). Additionally, analysis of the overall ligand-receptor interactions showed higher CCL and CXCL signaling pathways in sSDN4 environment compared with sSDN2 (Figure S7B). The signal directionality indicated that sSDN2 neutrophils received more signals from surrounding cells compared to sSDN4 neutrophils, which were predominantly signal senders (Figures S7C and S7D). Specifically, VTN–PLAUR interaction were found between tumor cells and neutrophils in sSDN2, which may play a role in the reprogramming neutrophils via cell adhesion and inhibition of efferocytosis^40^ (Figure S7E). In contrast, sSDN4 showed increased fibroblast–neutrophil CXCL12–CXCR4 signaling and neutrophils–TEC CXCL8–ACKR1 signaling, which may enhance neutrophil recruitment to inflamed tissues^41,42^ (Figure S7E).

To validate the distinct neutrophil molecular profiles and cellular interactions associated with the two sSDNs, we generated single cell spatial transcriptome data from all tumor samples in the TIGER-LC cohort using the CosMx platform with a 6K gene panel. Analysis of this dataset confirmed the neutrophil gene expression patterns and cell–cell interaction patterns observed in the scRNA-seq data (Figures S8A-S8D).

To test whether tumor cell proximity in sSDN2 contribute to neutrophil reprogramming, we performed *in vitro* co-culture experiments using human peripheral blood neutrophils and two CCA cell lines, HuCCT1 and HuH28, with HEK293T cells as a control. We assessed *MMP9* and *TGFB1* expression as readouts of pro-tumor neutrophil polarization^43–46^. Compared with control conditions (neutrophils cultured alone or co-cultured with HEK293T), neutrophils co-cultured with CCA cell lines showed significantly higher expression of both markers (Figures S8E and S8F).

Together, these results suggest two contrasting neutrophil-associated microenvironments in iCCA exhibiting distinct functional profiles. One was a tumor-enriched environment in which MPO+CD15+ neutrophils co-localized with tumor cells and immunosuppressive cells. The other was a tumor-desert environment where MPO+CD15+ neutrophils co-localized with mostly immune-activating cells. These two environments were enriched in C4 and C3, respectively, which were associated with distinct patient outcomes.

### Signatures of sSDN2- and sSND4-associated neutrophil environments predict patient prognosis

To transfer the findings from spatial analysis to bulk transcriptome data, we developed a scoring system using sSDN2- and sSND4- neutrophils associated environments to stratify iCCA patients. Specifically, we generated two gene marker sets by leveraging the enriched cell types in sSDN2 and sSDN4 neutrophil surroundings (Figures 5G, 5H, and 6A). The sSDN2 score was calculated based on marker genes for MPO+CD15+ neutrophils (*MPO, FUT4*), tumor cells (*KRT17, KRT19, EPCAM)*, CD68-CD163+ Mo/Mac cells (*CD163*), Tregs (*FOXP3*), and CD14+ monocytes (*CD14*). The sSDN4 score was based on MPO+CD15+ neutrophils (*MPO, FUT4*), CD68+CD163+/- TAMs (*CD68*), cytotoxic T/NK cells (*GZMB, B3GAT1, CD8A, CD8B*), and CD38+ leukocytes (*CD38*). Since sSDN4 neutrophil surroundings had an absence of tumor cells, we further divided the sSDN4 score by average tumor cell gene expression (*KRT17, KRT19, EPCAM*) as the final sSDN4 score. CD11c+ DC/monocytes were not included in the scoring system because of the low enrichment compared with other cell types. Patients were then stratified based on median cutoff of each sSDN within each cohort and further categorized into three groups: sSDN2^high^sSDN4^low^, sSDN2^low^sSDN4^high^, and an intermediate “Other” group. We applied the scoring system in the TIGER-LC cohort and two additional independent iCCA cohorts, the International Cancer Genomics Consortium (ICGC) and Japan cohorts with 81 and 162 patients, respectively. Across all cohorts, the sSDN2^low^sSDN4^high^ group exhibited the most favorable overall survival, whereas the sSDN2^high^sSDN4^low^ group had the poorest prognosis (Figure 6B). Patients in the intermediate group showed moderate outcomes (Figure 6B). Notably, these survival associations remained highly consistent in both high- and low-tumor-purity patient groups (Figures S9A and S9B). Multivariate analysis of features from each cell type revealed inconsistent association with patient outcomes across cohorts (Figure S9C). In contrast, sSDN-based scoring system consistently achieved strong and reproducible survival prediction across all cohorts (Figure S9C). Taken together, these results demonstrate that neutrophil-associated spatial environments captured by sSDN2 and sSDN4 can be translated into robust gene signatures for prognostic stratification across independent iCCA cohorts.

**Figure 6.**
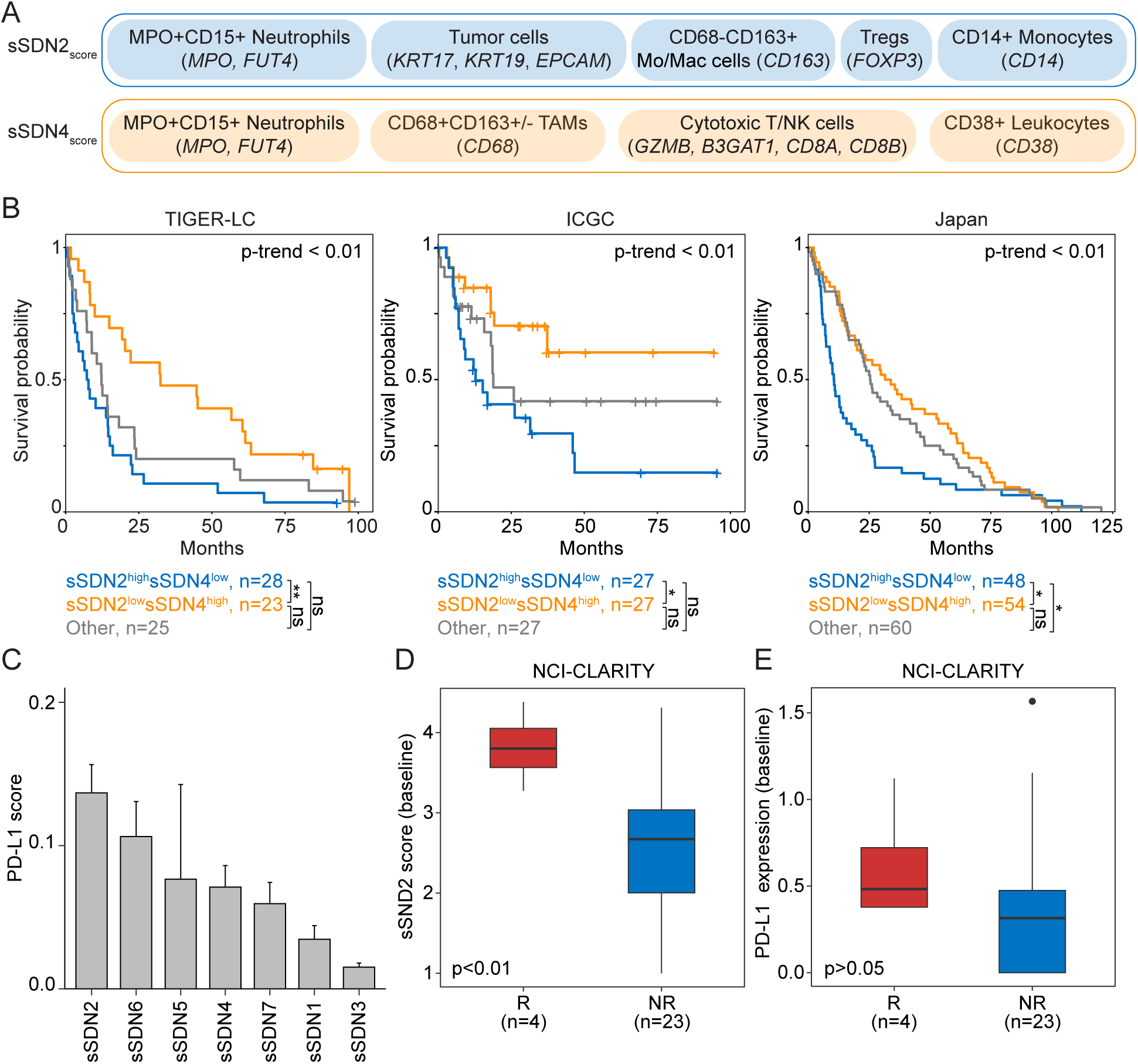
Features from MPO+CD15+ neutrophil-associated environments stratify patient prognosis in iCCA. (A) Illustration of the features used to represent sSDN2 and sSDN4 MPO+CD15+ neutrophil-associated environments. (B) Kaplan-Meier plots showing the overall survival of the iCCA patient groups in the TIGER-LC (left), ICGC (middle), and Japan (right) cohorts. *p*-value was calculated using log rank test and *p*-trend was calculated using log rank test for trend. \**p<*0.05, \*\**p*<0.01, ns, not significant. (C) PD-L1 scores of each sSDN. The PD-L1 score was calculated as the average PD-L1 expression of each sSDN in individual patients, normalized to tumor cell content. (D) Baseline sSDN2 scores in responders (R, n=4) and non-responders (NR, n=23) from the NCI-CLARITY cohort. (E) Baseline PD-L1 expression level in responders (R, n=4) and non-responders (NR, n=23) from the NCI-CLARITY cohort.

Consistent with the immunosuppressive role of sSDN2 in iCCA, we found that sSDN2 had the highest PD-L1 score among all the sSDNs (Figure 6C). We then investigated whether iCCA enriching for sSDN2 is responsive to checkpoint inhibitor-based immunotherapy. We calculated sSDN2 scores in baseline tumor samples collected from iCCA patients enrolled in immunotherapy clinical trials from our NCI-CLARITY study^47^. Consistently, patients who responded to immunotherapy (complete or partial response) had significantly higher sSDN2 scores compared to non-responders (stable or progressive disease; Figure 6D). In contrast, baseline PD-L1 expression levels only showed borderline, not statistically significant difference between responders and non-responders (Figure 6E). These findings suggest that immunotherapy may relieve the immunosuppressive environment and the sSDN2 score may serve as an effective predictor of immunotherapy response in iCCA. Given the limited number of patients in our cohort, validation in larger iCCA patient cohorts would further establish its predictive value.

## Discussion

The spatial organization of tumors is increasingly recognized as a key determinant of cancer biology, yet remains challenging to define due to the complexity and heterogeneity of cellular interactions within the TME. Here, we resolve tumor spatial architecture at single-cell resolution by modeling cell type-centric local multicellular interaction networks and their higher-order spatial organization. Using a large-scale spatial omics dataset comprising approximately 7 million cells from 162 iCCA patients, we defined 41 cell type-centric SDNs that captured the immediate cellular neighborhoods of individual cell types. Integration of these local networks revealed seven higher-order spatial environments (sSDNs), enabling multiscale characterization of tumor architecture. Different from global neighborhood learning approahces^17,48,49^, which often treat spatial organization as an aggregate property of cellular composition, this study explicitly captures the local cell-centered interaction networks as elementary units of spatial organization and reconstructs higher-order architecture through their integration. This strategy reveals structured relationships among these local networks within higher-order spatial contexts that are not captured by bulk or global neighborhood analyses. For example, tumor-enriched regions are not homogenous, but instead comprise multiple distinct interaction networks, including tumor cell-dominant, tumor-stromal, and diverse tumor-immune configurations. Moreover, interactions among these networks within higher-order environments are structured rather than random, indicating an additional layer of spatial organization. Importantly, the identified SDN and sSDN structures are reproducible across independent cohorts, spatial modalities and tissue scales, and consistently associated with patient outcomes, supporting the biological relevance and generalizability of the defined spatial architectures. Together, these cell type-centric SDNs and higher-order sSDNs define the spatial landscape of iCCA, demonstrating its utility in resolving complex spatial ecosystems.

Increasing evidence suggests that the spatial location of immune cells is a critical determinant of their functional roles in tumor progression and patient outcomes^1,2,50^. While prior studies have linked neutrophil infiltration to poor prognosis in iCCA, these analyses largely lack spatial resolution^10,51–53^. In this study, we identified two distinct neutrophil-associated microenvironments with opposing clinical outcomes by using our spatial framework. Neutrophil-associated tumor-enriched environments were related to immunosuppressive features, whereas neutrophil-associated tumor-desert environments were linked to immune-activating contexts. These spatially defined neutrophil populations showed distinct transcriptional profiles and interaction networks, suggesting functional heterogeneity shaped by spatial context. Although further experiments are required to establish causality, these findings highlight the importance of spatial context in shaping immune cell function within tumors.

Collectively, this study establishes a multilevel framework for defining tumor spatial architecture and resolves the complex spatial ecosystem of iCCA. By defining cell type-centric local networks and their higher-order organization, this approach enables systematic interrogation of spatial biology. Integration of large-scale spatial datasets will be essential for uncovering context-dependent cellular functions and may inform the development of spatially guided therapeutic strategies to improve cancer patient care. Beyond iCCA, this work also presents one of the largest single-cell spatial atlases in this disease context and provides a valuable resource for exploring multicellular interactions and tumor spatial organization.

## Methods

### Human sample collection

The TIGER-LC cohort comprised tumor and adjacent non-tumor paired samples collected from 131 patients with an initial diagnosis of iCCA. All patients gave informed consent with approval by Institutional Review Board (IRB). The clinical characteristics of these patients are provided in Table S1. All tumor (n=131) and non-tumor (n=131) samples were stained using CODEX. All tumor samples (n=131) were used for CosMx. Paired samples from a subset of 91 patients were subjected to bulk transcriptome profiling in our previous study^11^. Note that twelve patients were later re-evaluated due to potential errors in medical records, of which ten were confirmed by pathology to have cancer and two were excluded due to inappropriate documentation. The European iCCA patient cohort comprised 42 tumor samples from 31 patients (Table S3). All samples were subjected to whole-tissue slide-based CosMx profiling. All patients provided informed consent with IRB approval. Human peripheral blood was obtained from healthy donors enrolled under an IRB-approved NIH protocol (99-CC-0168).

### TMA construction for the TIGER-LC cohort

TMA was constructed for tumor and paired non-tumor tissues from 131 patients in the TIGER-LC cohort. Specifically, 1.0-millimeter (mm) cores from formalin-fixed paraffin-embedded (FFPE) tissues were assembled by the pathologist team from Laboratory of Pathology, NCI. Hematoxylin and eosin (H&E) staining was performed for slide sections for quality control.

### Antibody-conjugation with oligonucleotide-tags

Carrier free antibodies were conjugated to selected barcodes using commercial reagents following Akoya Bioscience’s recommended protocols as described earlier^16^. Briefly, preservatives and other additives (e.g., trehalose) were removed with Amicon Ultra 30K centrifugation filters (Millipore, Cat. No. UFC503024). Antibodies were washed multiple times with PBS, and the protein concentration was determined using an Implen nanophotometer set to mouse IgG parameters. For conjugation, 50 µg of protein was diluted in 100 µL PBS. One Amicon Ultra 50K centrifugation filter (Millipore, Cat. No. UFC505024) was used for each conjugation. Each filter was washed with 500 µL filter blocking solution (Akoya Biosciences, Cat. No. 7000009 Part No 232113) and centrifuged at 12,000 x g for 2 min. The remaining solution was discarded. Subsequently, 50 µg antibody in a volume of 100 µL supplemented with PBS was added to the filter and centrifuged at 12,000 x g for 8 min. 260 µL antibody reduction master mix (Akoya Biosciences, Cat. No. 7000009 Part No. 232114 and Part No.232115) was added to the filter and incubated for 30 min. Next, the filter was centrifuged at 12,000 x g for 8 min and washed with 450 µL of conjugation buffer (Akoya Biosciences, Cat. No. 7000009 Part No. 232116). The barcode (one vial for each antibody) was resuspended in 10 µL nuclease free water (Ambion AM9938) and mixed with 210 µL of conjugation buffer. The barcode solution was added to the filter and incubated for 2 hours. The filter was centrifuged at 12,000 x g for 8 min, washed thrice with 450 µL purification solution (Akoya Biosciences, Cat. No. 7000009 Part No. 232117) and resuspended into 100 µL of antibody storage solution (Akoya Biosciences, Cat. No. 7000009 Part No. 232118). To collect the conjugated antibody, the filter was inverted and centrifuged into a new tube at 3000 x g for 2 min. The collected conjugated antibody was stored at 4°C.

### CODEX staining

TMA slides of 4 µm thick were stained using CODEX (Phenocycler Fusion) with commercially available and custom conjugated antibodies (51 markers with 2 additional nuclear markers; Table S2) following recommendations from Akoya Biosciences. Specifically, slides were baked overnight at 60°C to improve attachment of tissues to glass, immersed in xylene (2 times 10 min) to remove paraffin and rinsed in a decreasing alcohol series to rehydrate the tissues (100%, 100%, 90%, 70%, 50%, 30% ethanol in MilliQ water for 4 min each). Antigen retrieval was performed in a pressure cooker set to low pressure using antigen retrieval solution pH 9 for 15 min. The slides were then cooled down to room temperature and soaked in Hydration (2 times 2 min) and Staining buffer (up to 20 min). Antibodies were combined in staining buffer containing J, N, S and G blocking reagents and added to tissues overnight at 4°C. The slides were washed in staining buffer (2 times 2 min) and the bound antibodies were fixed using 1.6% paraformaldehyde (diluted in storage buffer) for 10 min. Next, the slides were washed in PBS (3 times 2 min), incubated in ice cold methanol for 5 min, additionally washed in PBS (3 times 2 min), and incubated in the crosslinker solution diluted in PBS for 20 min. After additional PBS washes (3 times 2 min), the stained slides were imaged on Phenocycler fusion or stored at 4°C in storage buffer.

### CODEX imaging and image preprocessing

A flowcell was created for the slide and the imaging run was set-up according to the vendor’s recommendations. The generated images were uploaded into HALO image analysis software (NCI-HALO version 3.6) for visualization and data preprocessing. Nuclear segmentation based on DRAQ5 was done using the HALO Image Analysis Platform from Indica labs using the HALO AI v4.0 module. The marker intensity positivity was manually tuned for each of the 51 markers and inspected on HALO. For each protein marker, the same gating threshold was applied across all samples in each TMA. All intensities below the marker thresholds were set to 0 before quality control of cells. We excluded TMA cores from 15 patients with no DAPI staining during the initial quality control of images.

### CODEX data quality control and cell clustering

We adopted an unsupervised clustering method^54^ to annotate the phenotypes of 600,720 segmented cells from tumor samples with all 51 non-nuclear staining markers. For quality control, we removed cells with no detected marker expression and removed features expressed by less than 3 cells. This step resulted in 544,131 cells and all 51 markers. Since the tumor samples were imaged in two batches, the expression data was scaled by batch with *ScaleData* in R package Seurat (v4.4.0), reduced in dimension with principal component analysis (PCA) and integrated with Harmony^55^ (*RunHarmony* function in Seurat) based on the top 20 principal components (PCs). The top 40 harmony embeddings were selected for Louvain clustering using *FindNeighbors* and *FindClusters* functions (resolution=2) and visualized with Uniform Manifold Approximation and Projection (UMAP). Cluster annotation was done in a hierarchical approach. Clusters with 1 marker expression were annotated by the marker. The remaining clusters were first annotated with coarse categories: b-Catenin+E-Cadherin+CD45- (epithelial cells), CD45+b-Catenin-E-Cadherin- (immune cells), CD31+/CD34+ (endothelial cells), αSMA+ (fibroblasts). Then, the clusters were annotated based on additional markers. For example, HNF4α+ epithelial cells were annotated as HNF4α+ epithelial cells or hepatocytes and EpCAM+/PanCK+ epithelial cells were annotated as tumor cells or biliary epithelial cells depending on the tissue origin (tumor or non-tumor tissue) and confirmed using histology images. Remaining cells expressing non-discriminative marker(s), such as CD141 only or CD39 only were labeled as “Unknown.” We identified 6 main cell types (epithelial/tumor cells, endothelial cells, fibroblasts, lymphocytes, myeloid cells, unknown/other) and 41 cell subtypes. Immune cells include lymphocytes, myeloid cells, and other immune cells (CD45+ only).

The same steps were done for the non-tumor samples with 577,473 segmented cells and 43 markers after quality control (Louvain clustering, 30 PCs, 0.8 resolution). We identified the 5 main cell types (epithelial/tumor cells, endothelial cells, lymphocytes, myeloid cells, unknown/other) and 22 cell subtypes.

### SDN identification

We constructed a GAT to identify multicellular spatial organizations of cells across samples^56^. The GAT model takes both an adjacency list describing cell-cell neighboring information and a cell feature matrix as inputs. It then dynamically aggregates information of each cell’s neighbors into a learned embedding matrix used for downstream analysis.

#### Undirected neighboring graph and adjacency list construction

We used Giotto R package (v1.1.2) *createSpatialNetwork* function (method=Delaunay, k=20, maximum_distance_delaunay=40 µm) to construct a graph in which nodes represent cell centroids and an edge links two cells if they are within 40 µm (100 px) Euclidean distance away. The graph was then converted to an adjacency list and each edge was duplicated in both directions to represent an undirected graph. An adjacency list was constructed for each sample and merged. Cells across all samples had unique cell indices to prevent connections between samples.

#### Cell feature matrix construction

Each cell is represented by a merged one-hot encoding of its cell type and cell subtype annotations. For tumor cells, we merged all subtypes as “Tumor cells” to avoid the learned spatial structures driven mainly by tumor subtype differences and reduce complexity introduced to the model. For non-tumor cells, all main cell types and subtypes were used.

#### Graph attention networks

The GAT was built with GATv2Conv module implemented in PyTorch (v1.21.1) and has 128-dimensional hidden representation, 3 layers to aggregate neighbor information within 3 hops away from the root cell (0-hop), and 32-dimensional output embeddings. Self-attention loops were removed to maximize focus on neighbor cells. We referred to Shiao et al. for GAT training^48^. During training, an immediate neighbor and a random cell is sampled for each target node. The loss function assumes that the immediate neighbor should be more similar to the target cell than a randomly sampled cell. For training, we ran 150 epochs, 5e-3 learning rate with Adam optimization algorithm, learning rate gamma of 0.1, and 2 attention heads. We trained with a batch size of 3096 cells randomly sampled from input.

#### Embedding clustering and SDN annotation

To identify cell type-centric SDNs, we split the GAT output embedding matrix based on the root cell type (tumor/epithelial cells, TECs, CAFs, lymphocytes, and myeloid cells). For each cell type, we ran PCA, *FindNeighbors, FindClusters,* and Louvain clustering (top 31 PCs, res=0.2) on the embeddings. Then, for each cluster, we computed the cell type composition of all immediate (1-hop) neighbors according to the neighboring graph. The clusters were annotated based on this aggregated neighbor cell type composition, excluding the root cells. We obtained 8 SDN_Ts for tumor/epithelial cells, 9 SDN_Es for TECs, 6 SDN_Fs for CAFs, 8 SDN_Ls for lymphocytes, and 10 SDN_Ms for myeloid cells. All SDNs labeled with “HNF4α+Ep” contained high proportions of HNF4α+ epithelial cells lacking tumor markers.

### sSDN characterization

To characterize high-order cellular organizations, we ran the same GAT network described above to learn neighboring SDN interactions. The feature matrix was constructed with all SDN annotations instead of cell types. To capture higher-order cellular organizations than SDNs, GAT model was expanded to 4 layers for 4-hop neighborhoods. Note that the maximum distance between 1-hop neighbors is 40 µm Euclidean distance as before. Finally, we trained with 1 attention head with all other hyperparameters the same. Louvain clustering with all 31 PCs and 0.1 resolution yielded 10 clusters. Two clusters highly represented in less than three samples were removed from downstream analysis. Additionally, one cluster enriched in epithelial cells lacking tumor markers were excluded. We obtained 7 final sSDNs.

### Tumor subtype SDN enrichment

To evaluate whether each tumor subtype preferred specific SDN environments, the tumor subtypes were randomly permutated across all tumor cells 1000 times. Enrichment was calculated as the log2 fold change of the observed-over-permutated proportions of tumor subtypes surrounded by each SDN. Two-sided *p*-values were calculated as the number of times observed was greater or less than permutated and adjusted by FDR.

### Neighbor cell type enrichment

To evaluate whether a cell type co-localize with specific cell types, all cell types of interest were randomly permutated across 1000 times. Enrichment was calculated as the log2 fold change of the observed-over-permutated cell type proportions within a 40 µm. Two-sided *p*-values were calculated as the number of times observed was greater or less than permutated and adjusted by FDR.

### SDN-SDN co-localization within each sSDN

To determine the unique spatial cellular organization of each sSDN, we computed the enrichment of SDN-SDN co-localization patterns^57^. We ran random permutation (n=1000) of the cell SDN labels within the neighboring graph across all samples. The mean log2 fold change of observed-over-permutated frequencies of edges between two SDNs was computed as the interaction enrichment. Two-sided *p*-values were calculated as the number of times observed was greater or less than permutated frequencies and adjusted by FDR. A simulation was run for each sSDN. To minimize the uncertainty of these interactions, interactions containing SDNs with less than 0.05 proportions in a sSDN were removed.

### Statistical significance of the sSDN structures

Statistical significance of each sSDN structure was calculated using the Hotelling’s *T*-squared test between SDN composition of each structure compared with the SDN composition in each sample. For sSDN5 composed of 1 SDN, *p*-value was calculated with two-sided Student’s *t*-test.

### Immune cell enrichment in MPO+CD15+ neutrophil surroundings

The enrichment of immune cells in MPO+CD15+ neutrophil surroundings were computed in a 40 µm radius area centered around MPO+CD15+ neutrophils selected in the following way. The sSDN2 MPO+CD15+ neutrophils were in sSDN2 and within 40 µm of a tumor cell. The sSDN4 MPO+CD15+ neutrophils were in sSDN4 and greater than 80 µm away from a sSDN2 cell to minimize surrounding overlap. The density of an immune cell subtype was computed as its frequency in a 40 µm radius of an MPO+CD15+ neutrophil divided by the area and averaged across all target MPO+CD15+ neutrophils. We ran random permutation (n=1000) of immune cell subtype labels. Enrichment score was the mean log2 fold change of observed-over-permutated densities and two-sided *p*-values were calculated as the number of times observed was greater or less than permutated densities and adjusted by FDR.

### Gene Set Enrichment Analysis (GSEA)

Differential expressed genes (DEGs) were first calculated using two-sided Student’s *t-*test. GSEA analysis was performed with R package fgsea (v1.32.4) on DEGs with a significance *p*<0.05 and an absolute log2 fold change greater than 0.01. Pathways with *p*-values <0.05 were considered as enriched pathways.

### Tumor purity

ESTIMATE^58^ was used to calculate tumor purity score for bulk transcriptome data.

### MaxFuse integration

The scRNA-seq data of 28 iCCA tumor samples from Xue et al.^10^ was filtered with 100 minimal features, clustered with Louvain clustering and annotated into neutrophils, TAMs, T cells, B cells, CAFs, TECs by following marker genes in the original study. The remaining clusters expressing *EPCAM, KRT17,* and *KRT19* were annotated as tumor cells. These 7 cell types were used during integration. MaxFuse^34^ was then applied to integrate the scRNA-seq data and the CODEX data. This method has been successfully used in previous studies to integrate datasets of weakly linked modalities, including proteomics and transcriptomics^59^. Specifically, a shared embedding was created iteratively using both common and distinct features between the two datasets. From the shared embedding, each CODEX cell was matched to a single cell from the scRNA-seq data. Then, the matches were filtered with score > 0.6, resulting in an average of 0.83 cell type annotation accuracy. As such, the mapped scRNA-seq cells could be grouped as sSDN2 and sSDN4 neutrophils, tumor cells, TECs, etc. To obtain neutrophil-specific genes associated with sSDN2 and sSDN4 separately, we first generated neutrophil-related genes by comparing genes from neutrophils with all other cell types (log2 fold change > 0.25, mean expression in neutrophils > 0.5, adjusted *p*-value < 0.05). We further performed differential gene expression analysis between sSDN2 and sSDN4 neutrophils (adjusted *p*-value < 0.05), followed by gene filtering using the neutrophil-related genes. Ligand-receptor interaction analysis between neutrophils and other cell types were performed using CellChat (v2.2.0)^60^.

### Scoring system for bulk transcriptome data

To translate spatial features to clinical scoring with bulk transcriptome data, the iCCA patients in each cohort were split into three groups based on the sSDN2 and sSDN4 neutrophil surrounding scores. We used gene markers representative of CODEX cell types enriched in each neutrophil surrounding environment. Tumor cells were represented by *KRT17* (PanCK), *KRT19* (PanCK), and *EPCAM*; MPO+CD15+ neutrophils by *MPO* and *FUT4* (CD15); CD163+ Mo/Mac by *CD163*; CD14+ monocytes by *CD14*; Tregs by *FOXP3*; CD68+ TAMs by *CD68*; cytotoxic T/NK or NK cells by *GZMB*, *B3GAT1* (CD57), *CD8A* (CD8), and *CD8B* (CD8); and CD38+ leukocytes by *CD38*. Since a family of KRT genes encode the PanCK protein, we computed DEGs between tumor and non-tumor samples using the paired bulk transcriptome data and found *KRT17* and *KRT19* to be enriched in tumor samples with higher-than-average overall expression. The expressions of these genes were used to calculate scores for each patient:

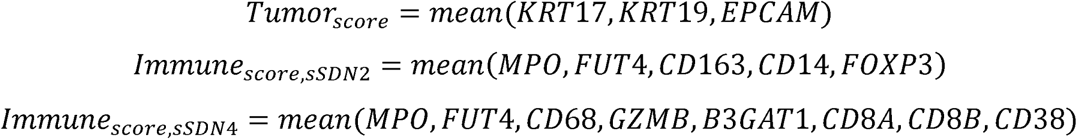

Since the sSDN2 neutrophil surroundings were enriched in tumor cells and immune cells, we maximize both with the following formula:

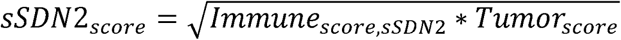

Since the sSDN4 neutrophil surroundings were enriched in immune cells but depleted in tumor cells, we maximize immune and minimize tumor with the following formula:

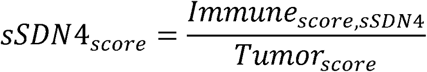

Finally, patients were split into two groups based on the medians of *sSDN2_score_* and *sSDN4_score_* within each patient cohort and further combined into “sSDN2^high^sSDN4^low^”, “sSDN2^low^sSDN4^high^”, and the remaining as “Other.” We applied the same stratifying criteria for all cohorts.

### Overall survival

The Kaplan-Meier plot was applied for overall survival analysis using the R package Survival (v3.8-3).

### CosMx SMI sample preparation

Tissue samples were prepared for CosMx SMI according to He et al.^15^. In brief, 5-μm FFPE tissue sections were mounted onto VWR Superfrost Plus slides and baked overnight at 60 °C to ensure tissue adherence. Following the deparaffinization and target retrieval protocol, slides were fitted with an incubation frame (CosMx FFPE Slide Prep Kit RNA). Protein digestion was carried out using 3 μg/mL Proteinase K (Bruker) at 40 °C for 30 min. Tissues were incubated in 1:400 diluted fiducials (Bangs Laboratories) in 2× saline sodium citrate (SSC) and 0.001% Tween-20 (Teknova) for 5 min at room temperature, then washed in 1× PBS (ThermoFisher) for 5 min. Sections were fixed with 10% neutral-buffered formalin for 1 minute at room temperature, washed according to protocol, and blocked with 100 mM *N*-succinimidyl (acetylthio) acetate (NHS-acetate, ThermoFisher) in NHS-acetate buffer (0.1 M sodium phosphate, 0.1% Tween-20, pH 8, in DEPC H O) for 15 minutes at room temperature. Following a 5-minute rinse in 2X SSC, Bruker ISH probes were incubated at 95 °C for 2 minutes and cooled on ice. The hybridization mixture (1 nM ISH probe, 10 nM attenuation probes, 1X Buffer R, 0.1 U/μL SUPERase•In™ [ThermoFisher] in DEPC H O) was applied to the slides, which were sealed, hybridized at 37 °C for 16–18 hours, and washed. Finally, the slides were attached with custom-made flow cells.

### CosMx imaging

We applied CosMx 6K gene panel for TIGER-LC cohort and four samples (c1, c2, c3-2, c60-2) from the European cohort. The rest of the samples from the European cohort were profiled using CosMx 1K gene panel. RNA targets were detected using the CosMx SMI instrument following the procedure by He et al.¹. Briefly, tissue sample was assembled into the flow cell, mounted onto the SMI instrument, and washed with reporter wash buffer to remove air bubbles. Fields of views (FOVs) were aligned with the entire tissue area. For RNA readout, the flow cell was washed with 100 μL of reporter buffer 1, incubated for 15 min, washed again with 1 mL reporter wash buffer, and finally washed with 100 μL imaging buffer. Each FOV was captured by eight Z-stack images at 0.8 μm intervals. Then, fluorophores attached to reporter probes were UV cleaved and washed with 200 μL strip wash buffer. This fluidic and imaging sequence was repeated for multiple reporter pools, and the entire cycle was repeated multiple times to increase RNA detection sensitivity. Lastly, tissue and membrane morphology were visualized using oligonucleotide-conjugated antibodies hybridized to barcode reporters. In this step, the flow cell was incubated with blocking buffer (buffer W, Bruker) for 30 min, incubated with four antibody (CD298, CD45, CD68, PanCK) and DAPI stains for 1 h, washed with 9 mL reporter wash, and then washed with 100 μL imaging buffer. Eight Z-stack images were acquired across five fluorescence channels (four protein markers and DAPI).

### CosMx data quality-control and cell type annotation

For the whole-tissue slide-based CosMx data in the European cohort, we retained 5,558,983 high-quality cells for downstream analysis after quality control (≥ 20 total counts and ≥ 10 genes detected in each cell). Because we used 1K gene panel for most of the samples in the cohort, we applied our previously published scRNA-seq dataset^5^ for label transfer to ensure the high accuracy for cell type annotation. Specifically, we identified three major cell types, i.e., epithelial cells, immune cells, and stromal cells, via Harmony (v1.2.0) label transfer based on the top 20 PCs, followed by cell clustering using the top 10 Harmony embeddings with default parameters. Clusters were annotated by the dominant scRNA-seq-derived cell type and marked as unclassified if the top cell type comprised < 50% of all or if the second-highest proportion was > 50% of the top. We further performed a second round of clustering within each major cell type (top 10 PCs, Louvain algorithm). Malignant cells and non-malignant epithelial cells were determined using DEGs combined with histology images as previously described^15^. Immune cells across samples were integrated using Harmony on the top 50 PCs and were annotated based on DEGs (adjusted *p*-value < 0.05, expression proportion ≥ 0.1, and log fold change > 0.5). Sixteen immune clusters were identified using the top 14 Harmony embeddings and default clustering parameters. Similar analyses were performed for stromal cells with 15 stromal clusters identified using the top 16 Harmony embeddings and default clustering parameters.

For the CosMx data from the TIGER-LC cohort, we performed quality control by keeping cells with ≥50 total counts and ≥10 genes detected within each cell. We further checked imaging quality of all FOVs across samples. A total of 272,823 high-quality cells were retained for downstream analysis. We performed PCA on 2,000 highly variable genes (HVGs) and ran clustering using the top 10 PCs (resolution = 0.5). Major cell types were annotated based on DEGs filtered by adjusted *p*-value < 0.05, expression proportion ≥ 0.1, and log fold change > 0.5. Identified cell types included malignant cells, epithelial cells, stromal cells, and immune cells. Immune cells were further classified into lymphocytes, macrophages, and neutrophils by clustering analysis using the top 10 PCs (resolution = 0.5) derived from PCA based on 1,000 HVGs across all immune cells.

### SDNs and sSDNs in the CosMx data

To map SDNs identified in the CODEX data to the CosMx datasets, we trained cell type-specific multi-class classification models using the XGBoost algorithm based on the CODEX data. To ensure compatibility between data modalities, models were trained using the major cell types (malignant, epithelial, stromal, macrophage, neutrophil, lymphocyte), given differences in cell type annotation strategies between CODEX (major lineage-marker based) and CosMx (unsupervised clustering-based). A 3-hop neighborhood of each target cell derived from Delaunay triangulation, along with the SDN assignment of the target cell from CODEX data were used in the training. Delaunay triangulation with 40 μm distance limit was computed using the R package deldir (v2.0-4). The dataset was randomly split into training (80%) and test (20%) sets, with 20% of the training data held out for surveillance to avoid model overfitting during the training. Models were built using the xgboost R package (v3.1.3.1), with objective = “multi:softprob” for probabilistic output and eval_metric = “mlogloss” to monitor performance. Regularization parameters included subsample = 0.8, colsample_bytree = 0.8, gamma = 1, and min_child_weight = 3. Models were trained for up to 1000 boosting rounds with early stopping after 20 rounds without improvement in validation loss. A watchlist was used to monitor both training and validation performance. Due to variability in the number and characteristics of cell type-specific SDNs, learning rate and maximum depth were optimized separately for each model: tumor cell SDNs (learning rate = 0.02, max depth = 7), TEC SDNs (learning rate = 0.015, max depth = 8), CAF SDNs (learning rate = 0.015, max depth = 5), lymphocyte SDNs (learning rate = 0.015, max depth = 7), and myeloid SDNs (0.02, max depth = 9). The trained models were then used to predict SDNs for CosMx datasets, the same 3-hop neighborhood features were used as input. Cells with predicted probabilities ≤ 0.5 were labeled as Unknown.

To map sSDNs, we trained a separate multi-class XGBoost model using similar pipeline and parameters with the following modifications. Features were derived from the major cell type composition within the 4-hop neighborhood of each target cell as used in sSDN identification in the CODEX data. All sSDNs used learning_rate = 0.1, and max_depth = 6. For predict sSDNs for the CosMx data, the same 4-hop neighborhood features were used as query dataset. To assess the prevalence of sSDNs across samples, we compared observed (o) to expected (e) cell numbers for each sSDN-sample combination, defined as Ro/e = observed/expected based on established method^61^. Expected cell numbers for each combination were calculated from Chi-square test.

### CosMx-based ligand-receptor analysis within sSDNs

We calculated ligand-receptor interaction scores using the spatially informed Hill function in CellChat (v2.2.0). Within each sample, trimmed mean (trim = 0.01) and ‘*computeRegionDistance*’ (interaction.range = 40, contact.range = 10, k.min = 10, tol = 5) were used to calculate gene expression and spatial distance. Samples with fewer than 100 cells in sSDN2 or sSDN4 were excluded to ensure robust estimation.

### Isolation and co-culture of neutrophils with cell lines

Neutrophils were isolated from whole blood obtained from healthy donors using the EasySep™ Direct Human Neutrophil Isolation Kit (STEMCELL Technologies,19666), according to the manufacturer’s instructions. Purified neutrophils were co-cultured with CCA cell lines (HuCCT1 and HuH28), HEK293T cells, or cultured alone using a Transwell system (0.4-µm pore size; 12-well plates). Neutrophils and CCA cell lines were maintained in RPMI-1640 medium (Thermo Fisher Scientific, 21870092), whereas HEK293T cells were cultured in DMEM (Thermo Fisher Scientific, 10313039). Both media were supplemented with 10% fetal bovine serum (Cytiva, SH30396), 100 U/mL penicillin, and 100 µg/mL streptomycin (Thermo Fisher Scientific, 15140122). Cells were incubated at 37 °C in a humidified atmosphere containing 5% CO. After 24 h of co-culture, neutrophils were harvested and processed for RT-qPCR analysis.

### RNA extraction, cDNA synthesis, and RT-qPCR

Total RNA was extracted from cell pellets using the RNeasy Mini Kit (Qiagen, 74104) according to the manufacturer’s instructions. Purified RNA was subsequently reverse-transcribed into cDNA using the High-Capacity cDNA Reverse Transcription Kit (Thermo Fisher Scientific, 4374966). Quantitative real-time PCR (RT-qPCR) was performed using SYBR™ Green Universal Master Mix (Thermo Fisher Scientific, 4364344) and gene-specific primers (Bio-Rad), following the manufacturer’s protocol. Relative gene expression levels were calculated using the comparative ΔΔCt method normalized to house-keeping control *GAPDH*.

## Data availability

CODEX data of the TIGER-LC cohort generated in this study are available at zenodo: https://doi.org/10.5281/zenodo.15419271. CosMx data of the European cohort and the TIGER-LC cohort generated in this study are available at zenodo: https://doi.org/10.5281/zenodo.18391527. The publicly available datasets used in this study include single-cell transcriptome data of <u>GSE151530</u>^5^ and <u>PRJCA007744</u>^10^, bulk transcriptomic data of <u>GSE76297</u> (TIGER-LC, iCCA)^11^, dbGaP: <u>phs003074.v1.p1</u> (NCI-CLARITY)^47^, <u>EGAS00001000950</u> (Japan), and <u>GSE89749</u> (ICGC)^62^.

## Acknowledgement

We thank Nicole Ducich and Qin Li for technical support, and the patients, families, and nurses for contribution to this study. This work was supported by grants (ZIA BC 012079, ZIA BC 012083, Z01 BC 010877, Z01 BC 010876, Z01 BC 010313 and ZIA BC 011870) from the Intramural Research Program of the Center for Cancer Research, National Cancer Institute of the United States. The contributions of the NIH authors were made as part of their official duties as NIH federal employees, are in compliance with agency policy requirements, and are considered Works of the United States Government. However, the findings and conclusions presented in this paper are those of the authors and do not necessarily reflect the views of the NIH or the U.S. Department of Health and Human Services.

## Author Contributions

L.M. developed the study concept. M.R. and J.U.M. directed the clinical study. H-P.L., M.L., and W.W. performed computational analysis. N.T.L.N., N.K., and M.O.H. conducted experiments. S.M.H. performed TMA design, construction, evaluation, and data interpretation. J.C., D.C., E.L., M.K., M.F., M-H.H., A.B., N.A., R.L., E.R., S.L., and X.W.W. performed additional analysis. H-P.L. and L.M. wrote the manuscript with help from M.L., W.W., and N.T.L.N. All authors read, edited, and approved the manuscript.

