## supplemental figures and tables for "Cell type-centric interaction networks define spatial architecture of intrahepatic cholangiocarcinoma"

### Supplementary Figure legends

**Figure S1. CODEX profiling of iCCA.**

(A) UMAP representation of 544,131 single cells expressing at least 2 markers from tumor samples after quality control. Colored and labeled by cell type.

(B) Cell (sub)type compositions for each tumor sample. Samples were hierarchically clustered. Refer to (A) for color coding.

(C) A representative tumor (LCS-639) showing CODEX image, cell type labels, and higher magnification of the CODEX staining for 2 regions with selected markers.

(D) UMAP representation of 577,473 single cells expressing at least 2 markers from non-tumor samples after quality control. Colored and labeled by cell type.

(E) Heatmap of the average marker expression level for cell (sub)types in non-tumor samples.

(F) Stacked bar plot showing the cell (sub)type composition in each non-tumor sample. Samples were hierarchically clustered. Refer to (E) for color coding.

(G) Bar plot of overall cell type proportions in tumor versus non-tumor samples. Biliary epithelial and tumor cells compare the proportion of biliary epithelial cells in non-tumor samples to that of tumor cells in tumor samples.

(H) Bar plot of immune cell subtype proportions in tumor versus non-tumor samples out of all cells in each tissue type. (G and H) Error bars represent standard error of the mean. For statistical tests, *p*-values were calculated using two-sided Student's *t*-test. *$p<0.05$, **$p<0.01$, ***$p<0.001$, ****$p<0.0001$.

**Figure S2. Cell type-centric SDNs.**

(A) UMAP of clustered tumor cell GAT embeddings labeled and colored by SDN_T.

(B) Heatmap of tumor cell subtype enrichment in each SDN_T. Enrichment values with FDR-adjusted *p*-values ≥ 0.05 were set to 0.

(C) Waterfall plot showing the log2 fold change (FC) of the mean number of neighbor tumor or immune cells within a 40 μm radius surrounding HLADR+ tumor cells over HLADR- tumor cells. FDR-adjusted *p*-value < 0.05.

(D) UMAP of clustered lymphocyte GAT embeddings labeled and colored by SDN_L.

(E-F) UMAP of clustered GAT embeddings colored by SDN_F (E) and SDN_E (F) clusters and heatmap of SDN_F (E) and SDN_E (F) cell subtype compositions. Cell subtypes with greater than 0.01 proportion are shown.

(G) A representative tumor sample shown with CODEX staining (left), SDN_E annotation (middle), and SDN_F annotation (right).

(H) UMAP of clustered myeloid cell GAT embeddings colored by SDN_M and heatmap of SDN_M cell subtype composition. Cell subtypes with greater than 0.01 proportion are shown.

(I) A representative tumor sample shown with CODEX staining (left) and SDN_M annotation (right).

(J) Bar plot showing the number of patients with each identified SDN. A red, dotted line marks y=3.

(E, F, and H) T/Ep, tumor cell/epithelial cell; CAF/TEC, cancer-associated fibroblast/tumor-associated endothelial cells; LyC, lymphocyte; MyC, myeloid cell; Neu, neutrophil; Leu, leukocyte; Fb, fibroblast; Unk, unknown.

**Figure S3. sSDN identification.**

(A) UMAP of sSDNs by clustering of learned GAT embeddings.

(B) Proportion of each cluster in (A) in individual tumor samples. Each dot represents a sample. Bar and whisker indicate the mean and standard error. Red dotted line at y=0.05 indicates that sSDN9 and sSDN10 make up less than 5% of most samples.

(C) Heatmap of SDN composition of each sSDN. SDNs with greater than 0.01 proportion across all sSDNs are shown.

(D and E) Bar plot showing the global Moran's I of each sSDN (D) and SDN (E) within individual samples. Only samples containing more than 50 cells of the corresponding sSDN or SDN were included. Bar and whisker represent the mean ± standard error of the mean.

(F and G) Proportions of sSDN2 and sSDN4 in patient groups C3 (F) or C4 (G). Each box shows the median (center line), interquartile range (box), and data range (whiskers). *p*-value was calculated with two-sided Student's *t*-test.

(H) Undirected neighboring graph of four samples.

**Figure S4. Characteristics of the four patient clusters determined by sSDNs.**

(A) Box plot comparing tumor cellularity across four patient groups. Boxes indicate the median (center line), interquartile range (box), and data range (whiskers). *p*-value was calculated using two-sided Student's *t*-test.

(B and C) Kaplan-Meier curves showing overall survival of four patient groups within the high (B) and low (C) tumor cellularity strata. Overall *p*-values were calculated using log-rank test.

(D) Hierarchical clustering of tumor samples by cell (sub)type proportions in each sample using Pearson correlation distance and "ward.D2" method. Tumor subtypes were combined as "Tumor cells." CC, composition-based clustering.

(E) Overall survival of the four iCCA patient groups in (D). *p*-value calculated using log rank test. CC, composition-based clustering.

(F) Overall survival of patients stratified by tumor-immune ratio, defined by the relative numbers of tumor and immune cells in each sample. *p*-value was calculated using the log-rank test.

(G) Waterfall plot showing the association between iCCA driver mutations and four patient groups. *p*-values were calculated using the chi-squared test and adjusted by FDR.

(H) Heatmap showing associations between the four patient groups and sSDN (top), SDN (middle), and clinical variables (bottom). Hierarchical clustering of sSDN and SDN was performed using Pearson correlation distance and "ward.D2" method. *p*-value was calculated using the chi-squared test and adjusted by FDR.

**Figure S5. Single-cell spatial transcriptome profiling of a European iCCA cohort.**

(A) Number of cells per sample from iCCA patients in the European cohort. Samples colored red were subjected to CosMx 6K-plex, and those colored black were subjected to 1K-plex gene panels.

(B) Expression of cell type-specific marker genes. Color indicates average gene expression and dot size stands for the fraction of cells expressing a certain gene.

(C) Stacked bar plot of cell type compositions in each tumor sample. Samples were hierarchically clustered.

(D) UMAP of immune cell clusters and heatmap of the average expression levels of differentially expressed genes in each cluster. Representative gene markers for each cell type are shown in the heatmap. Mph, macrophage; B, B cell; T, T cell; Neu, neutrophil; DC, dendritic cell.

(E) UMAP of stromal cell clusters and heatmap of the average expression levels of differentially expressed genes in each cluster. Representative gene markers for each cell type are shown in the heatmap. Fb, fibroblast; EC, endothelial cell; Mu, mural cell.

(F) The enrichment of SDNs within each sSDN. For each sSDN, "enriched-SDNs" represents the SDNs enriched in the corresponding sSDN in Figure 3A, while "non-enriched-SDNs" indicates the rest of the SDNs. Box plots showing the proportions of SDNs in the sSDNs. Each box shows the median (center line), interquartile range (box), and data range (whiskers). *p*-value was calculated with two-sided Wilcoxon test.

**Figure S6. Myeloid environments associated with sSDN2 and sSDN4.**

(A and B) Box plots comparing tumor cell proportions (A) and immune cell proportions (B) in C3 and C4. *p*-values were calculated with two-sided Student's *t*-test.

(C) Volcano plot of differentially expressed genes in C4 versus C3 patient clusters. Red indicates significantly upregulated genes in C4 with log2 fold change > 0.05 and blue indicates significantly downregulated genes in C4 with log2 fold change < -0.05.

(D) Enriched cancer hallmark pathways in C4 versus C3. X-axis represents normalized enrichment score (NES). FDR-adjusted *p*-values are indicated next to each bar. $^*p < 0.05$, $^{**}\ p < 0.01$, $^{***}\ p < 0.001$.

(E) Bubble plots of functional marker protein expression in immune cells enriched in sSDN2 (left) and sSDN4 (right) MPO+CD15+ neutrophil surroundings. Dot size indicates percent of cells expressing the marker and dot color indicates average protein expression aggregated at cell-level.

(F) Illustration of the method used to analyze spatial specificity of immune cells surrounding MPO+CD15+ neutrophils by computing immune cell densities in concentric rings of increasing radii. Density is calculated by dividing the frequency of an immune cell subtype in the ring by the ring area.

(G and H) Line plots showing the densities of the differentially enriched immune cell types in sSDN2 neutrophil (G) and sSDN4 neutrophil (H) surroundings. Y-axis represents the density normalized by the maximum density and x-axis represents the area marked by increasing radii. *Z* and *p*-values computed using the Mann-Kendall trend test shown for each trend.

**Figure S7. Analysis of neutrophils.**

(A) Volcano plot of neutrophil-related DEGs between neutrophils in sSDN2 versus sSDN4 using a scRNA-seq dataset. Red indicates significantly upregulated genes in neutrophils in sSDN2 with log2 fold change > 0.05, and blue indicates significantly downregulated genes in neutrophils in sSDN2 with log2 fold change < -0.05. Only neutrophil-related genes were included in this plot (Methods).

(B) Stacked bar plot of ranked relative ligand-receptor interaction pathway probability in sSDN2 and sSDN4, showing pathways with probability > 0.75. Color-coded by sSDN. X-axis labels colored by significant sSDN enrichment calculated using paired Wilcoxon test.

(C and D) Interaction weight and directions of ligand-receptor signals with neutrophils as source or target, indicated by arrow thickness and directions, in sSDN2 (C) and sSDN4 (D). Line thickness indicates interaction weight and color indicates cell type. Neu, neutrophil.

(E) Bubble heatmap showing the interaction probability of specific ligand-receptor pairs with neutrophils as either target (top) or source (bottom). X-axis represents cell type colored by sSDN origin. Interaction probability indicated by the size and color of each dot. Neu, neutrophil.

**Figure S8. Single-cell spatial transcriptome analysis of the TIGER-LC cohort.**

(A) UMAP of 272,823 single cells from tumor samples in TIGER-LC cohort resolved by CosMx after quality control. Cell types indicated by colors.

(B) Expression of cell type-specific marker genes. Color indicates average gene expression and dot size indicates the fraction of cells expressing a certain gene.

(C) Bar plot showing expression difference of the indicated genes in sSDN2 and sSDN4 neutrophils. *p*-values were calculated using a two-sided Wilcoxon rank-sum test.

(D) Ligand-receptor interactions score of indicated gene pairs in sSDN2 and sSDN4.

(E) Schematic overview of the experimental design for co-culture of peripheral blood neutrophils (PBNs) and CCA tumor or control cell lines.

(F) RT-qPCR of *MMP9* and *TGFB1* at 24h after PBN co-culture with CCA cell lines (huCCT1 and HuH28) or a control cell line (HEK293T). mRNA level was normalized to *GAPDH* and fold changed was calculated based on PBN culture alone. PBN, peripheral blood neutrophil. ANOVA test was used. **$p$<0.01, ***$p$<0.001, ****$p$<0.0001.

**Figure S9. Survival analysis using bulk transcriptome data.**

(A) Kaplan-Meier plots showing the overall survival of samples with high tumor purity scores split by groups shown in Figure 6B in the TIGER-LC (left), ICGC (middle), and Japan (right) cohorts. *p*-trend and pairwise *p*-values calculated using log rank test for trend and log rank test.
(B) Same as (A) but of samples with low tumor purity scores. *p*-trend and pairwise *p*-values calculated using log rank test for trend and log rank test.
(C) Hazard ratio plots of cell type multivariate cox model and sSDN2/sSDN4 based scoring system ($sSDN2^{high}sSDN4^{low}$ versus $sSDN2^{low}sSDN4^{high}$) in the TIGER-LC (left), ICGC (middle), and Japan (right) cohorts.

Figure S1

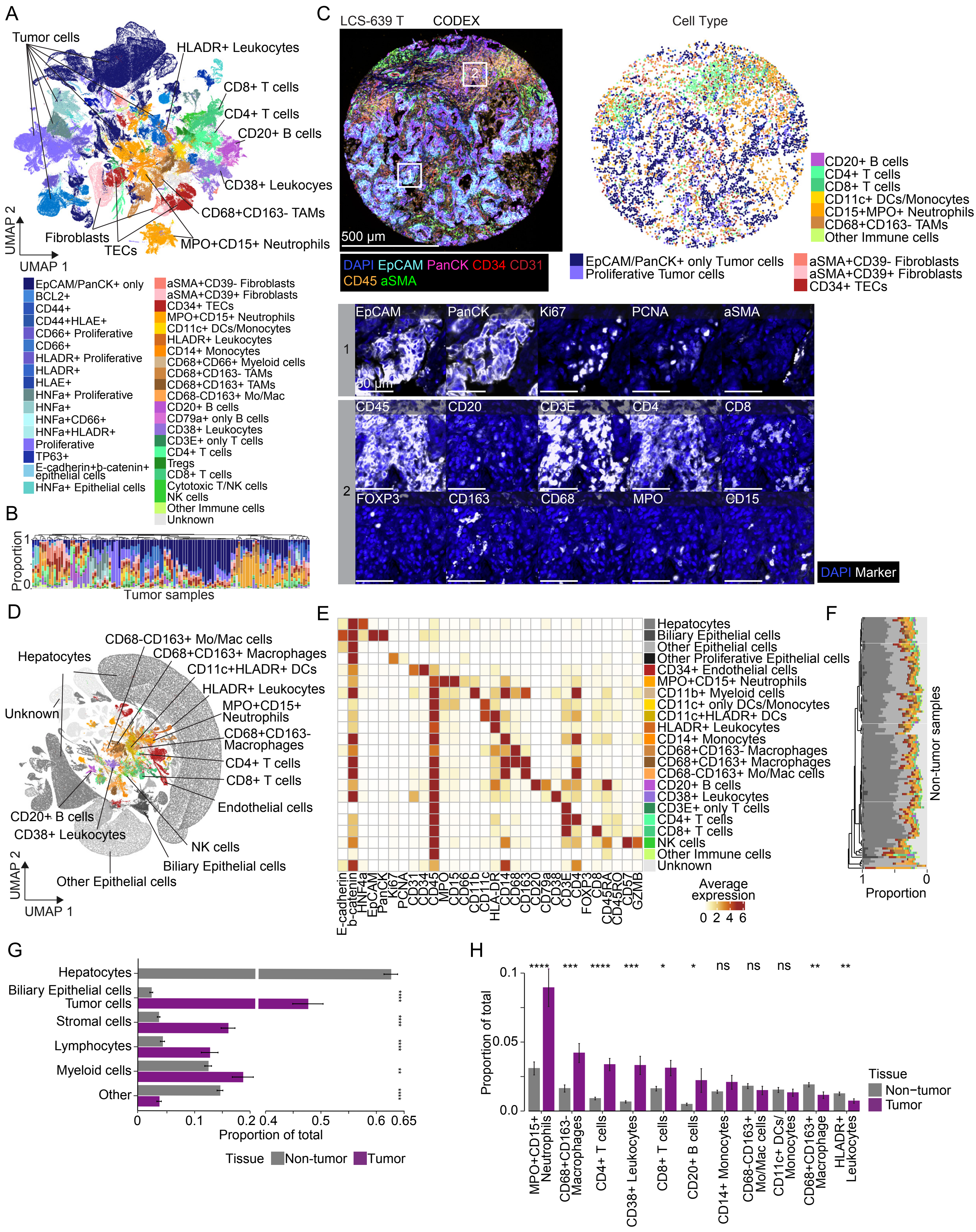
A
Tumor cells
HLADR+ Leukocytes
CD8+ T cells
CD4+ T cells
CD20+ B cells
CD38+ Leukocyes
CD68+CD163- TAMs
MPO+CD15+ Neutrophils
Fibroblasts
TECs
UMAP 2
UMAP 1
EpCAM/PanCK+ only
BCL2+
CD44+
CD44+HLAE+
CD66+ Proliferative
CD66+
HLADR+ Proliferative
HLADR+
HLAE+
HNFa+ Proliferative
HNFa+
HNFa+CD66+
HNFa+HLADR+
Proliferative
TP63+
E-cadherin+b-catenin+ epithelial cells
HNFa+ Epithelial cells
aSMA+CD39- Fibroblasts
aSMA+CD39+ Fibroblasts
CD34+ TECs
MPO+CD15+ Neutrophils
CD11c+ DCs/Monocytes
HLADR+ Leukocytes
CD14+ Monocytes
CD68+CD66+ Myeloid cells
CD68+CD163- TAMs
CD68+CD163+ TAMs
CD68-CD163+ Mo/Mac
CD20+ B cells
CD79a+ only B cells
CD38+ Leukocytes
CD3E+ only T cells
CD4+ T cells
Tregs
CD8+ T cells
Cytotoxic T/NK cells
NK cells
Other Immune cells
Unknown
B
Proportion
1
0
Tumor samples
C
LCS-639 T
CODEX
Cell Type
500 µm
DAPI EpCAM PanCK CD34 CD31
CD45 aSMA
CD20+ B cells
CD4+ T cells
CD8+ T cells
CD11c+ DCs/Monocytes
CD15+MPO+ Neutrophils
CD68+CD163- TAMs
Other Immune cells
EpCAM/PanCK+ only Tumor cells
Proliferative Tumor cells
aSMA+CD39- Fibroblasts
aSMA+CD39+ Fibroblasts
CD34+ TECs
1
2
EpCAM
PanCK
Ki67
PCNA
aSMA
50 µm
CD45
CD20
CD3E
CD4
CD8
FOXP3
CD163
CD68
MPO
CD15
DAPI Marker
D
CD68-CD163+ Mo/Mac cells
CD68+CD163+ Macrophages
CD11c+HLADR+ DCs
HLADR+ Leukocytes
MPO+CD15+ Neutrophils
CD68+CD163- Macrophages
CD4+ T cells
CD8+ T cells
Endothelial cells
Hepatocytes
Unknown
CD20+ B cells
CD38+ Leukocytes
NK cells
Biliary Epithelial cells
Other Epithelial cells
UMAP 2
UMAP 1
E
Hepatocytes
Biliary Epithelial cells
Other Epithelial cells
Other Proliferative Epithelial cells
CD34+ Endothelial cells
MPO+CD15+ Neutrophils
CD11b+ Myeloid cells
CD11c+ only DCs/Monocytes
CD11c+HLADR+ DCs
HLADR+ Leukocytes
CD14+ Monocytes
CD68+CD163- Macrophages
CD68+CD163+ Macrophages
CD68-CD163+ Mo/Mac cells
CD20+ B cells
CD38+ Leukocytes
CD3E+ only T cells
CD4+ T cells
CD8+ T cells
NK cells
Other Immune cells
Unknown
E-cadherin
b-catenin
HNF4a
EpCAM
PanCK
Ki67
PCNA
CD31
CD34
CD45
MPO
CD15
CD66
CD11b
CD11c
HLA-DR
CD14
CD68
CD163
CD20
CD79a
CD38
CD3E
CD4
FOXP3
CD8
CD45RA
CD45RO
CD57
GZMB
Average expression
0 2 4 6
F
Non-tumor samples
1
0
Proportion
G
Hepatocytes
Biliary Epithelial cells
Tumor cells
Stromal cells
Lymphocytes
Myeloid cells
Other
****
****
****
**
****
0
0.1
0.2
0.4
0.5
0.6
0.65
Proportion of total
Tissue
Non-tumor
Tumor
H
**** *** **** *** * * ns ns ns ** **
0.1
0.05
0.0
Proportion of total
MPO+CD15+ Neutrophils
CD68+CD163- Macrophages
CD4+ T cells
CD38+ Leukocytes
CD8+ T cells
CD20+ B cells
CD14+ Monocytes
CD68-CD163+ Mo/Mac cells
CD11c+ DCs/ Monocytes
CD68+CD163+ Macrophage
HLADR+ Leukocytes
Tissue
Non−tumor
Tumor

Figure S2

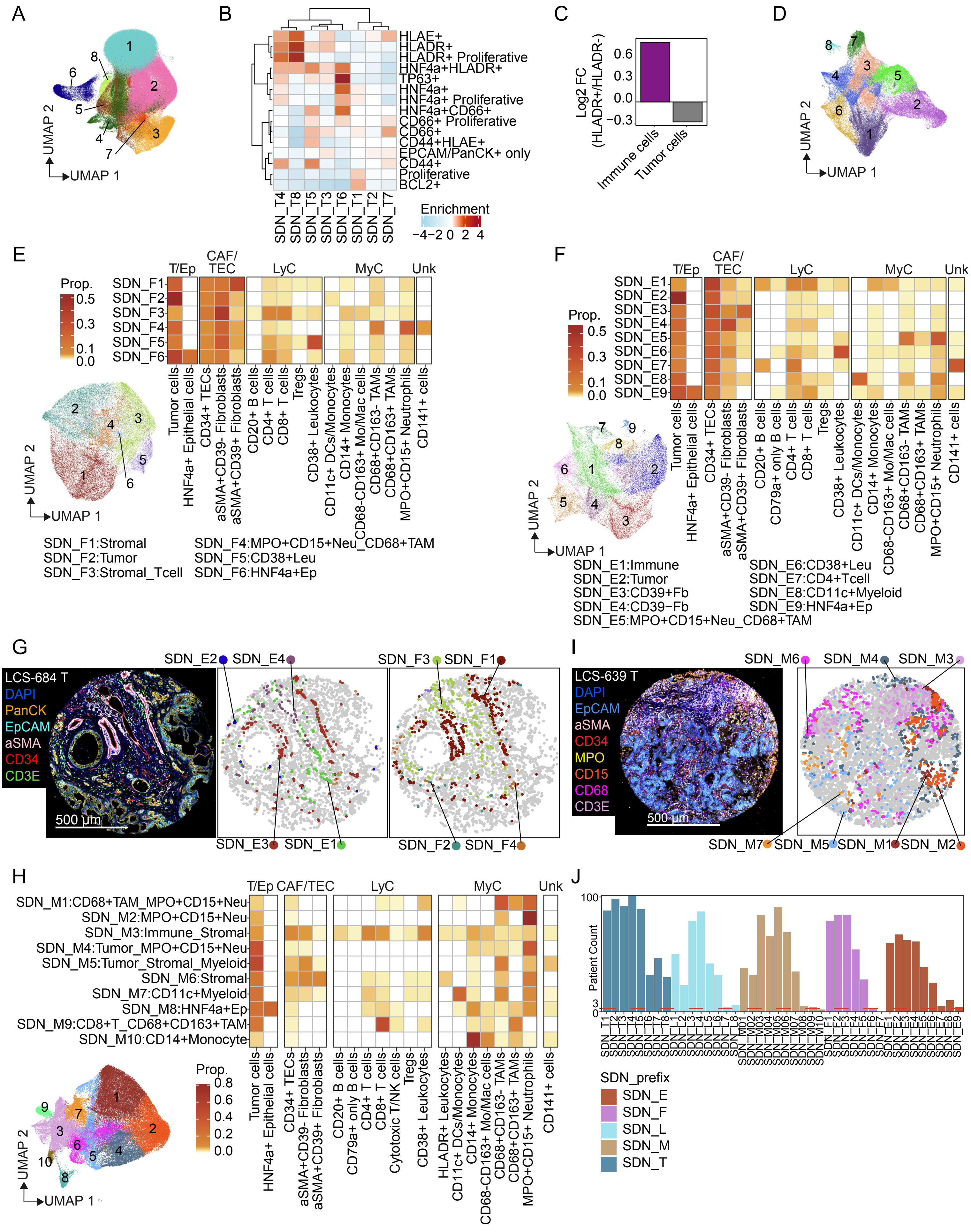
A
B
C
D
E
F
G
H
I
J
UMAP 1
UMAP 2
HLAE+
HLADR+
HLADR+ Proliferative
HNF4a+HLADR+
TP63+
HNF4a+
HNF4a+ Proliferative
HNF4a+CD66+
CD66+ Proliferative
CD66+
CD44+HLAE+
EPCAM/PanCK+ only
CD44+
Proliferative
BCL2+
SDN_T4 SDN_T8 SDN_T5 SDN_T3 SDN_T6 SDN_T1 SDN_T2 SDN_T7
Enrichment
−4 −2 0 2 4
Log2 FC (HLADR+/HLADR-)
Immune cells
Tumor cells
T/Ep
CAF/TEC
LyC
MyC
Unk
Prop.
Tumor cells
HNF4a+ Epithelial cells
CD34+ TECs
aSMA+CD39- Fibroblasts
aSMA+CD39+ Fibroblasts
CD20+ B cells
CD79a+ only B cells
CD4+ T cells
CD8+ T cells
Cytotoxic T/NK cells
Tregs
CD38+ Leukocytes
HLADR+ Leukocytes
CD11c+ DCs/Monocytes
CD14+ Monocytes
CD68-CD163+ Mo/Mac cells
CD68+CD163- TAMs
CD68+CD163+ TAMs
MPO+CD15+ Neutrophils
CD141+ cells
SDN_F1:Stromal
SDN_F2:Tumor
SDN_F3:Stromal_Tcell
SDN_F4:MPO+CD15+Neu_CD68+TAM
SDN_F5:CD38+Leu
SDN_F6:HNF4a+Ep
SDN_E1:Immune
SDN_E2:Tumor
SDN_E3:CD39+Fb
SDN_E4:CD39−Fb
SDN_E5:MPO+CD15+Neu_CD68+TAM
SDN_E6:CD38+Leu
SDN_E7:CD4+Tcell
SDN_E8:CD11c+Myeloid
SDN_E9:HNF4a+Ep
LCS-684 T
DAPI
PanCK
EpCAM
aSMA
CD34
CD3E
500 µm
LCS-639 T
MPO
CD15
CD68
SDN_M1:CD68+TAM_MPO+CD15+Neu
SDN_M2:MPO+CD15+Neu
SDN_M3:Immune_Stromal
SDN_M4:Tumor_MPO+CD15+Neu
SDN_M5:Tumor_Stromal_Myeloid
SDN_M6:Stromal
SDN_M7:CD11c+Myeloid
SDN_M8:HNF4a+Ep
SDN_M9:CD8+T_CD68+CD163+TAM
SDN_M10:CD14+Monocyte
Patient Count
SDN_prefix
SDN_E
SDN_F
SDN_L
SDN_M
SDN_T

Figure S3

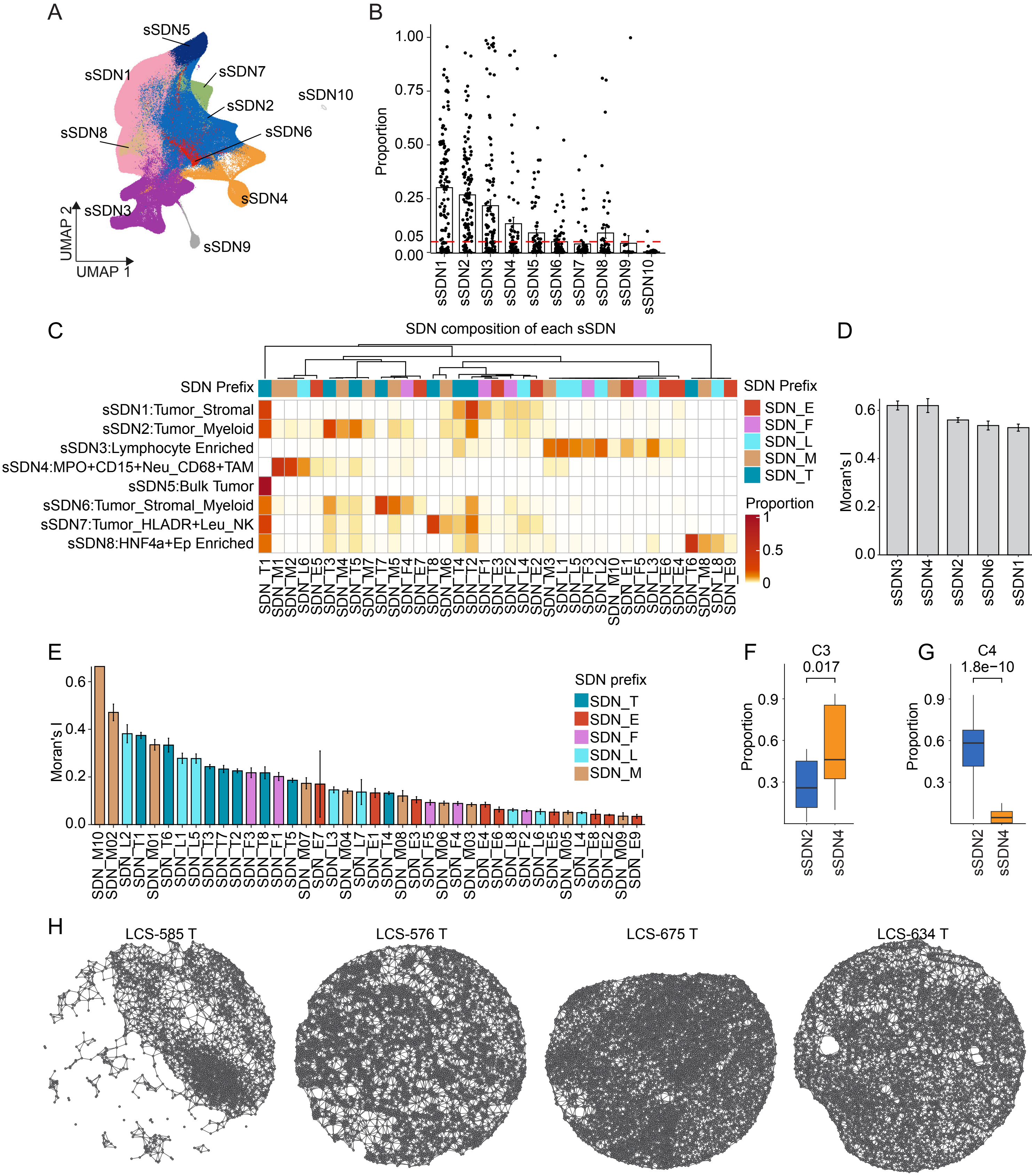

A
sSDN5
sSDN1
sSDN7
sSDN10
sSDN2
sSDN8
sSDN6
sSDN4
sSDN3
sSDN9
UMAP 2
UMAP 1
B
Proportion
1.00
0.75
0.50
0.25
0.05
0.00
sSDN1 sSDN2 sSDN3 sSDN4 sSDN5 sSDN6 sSDN7 sSDN8 sSDN9 sSDN10
C
SDN composition of each sSDN
SDN Prefix
sSDN1:Tumor_Stromal
sSDN2:Tumor_Myeloid
sSDN3:Lymphocyte Enriched
sSDN4:MPO+CD15+Neu_CD68+TAM
sSDN5:Bulk Tumor
sSDN6:Tumor_Stromal_Myeloid
sSDN7:Tumor_HLADR+Leu_NK
sSDN8:HNF4a+Ep Enriched
SDN_T1 SDN_M1 SDN_M2 SDN_L6 SDN_E5 SDN_T3 SDN_M4 SDN_T5 SDN_M7 SDN_T7 SDN_M5 SDN_F4 SDN_E7 SDN_T8 SDN_M6 SDN_T4 SDN_T2 SDN_F1 SDN_E3 SDN_F2 SDN_L4 SDN_E2 SDN_M3 SDN_L1 SDN_L5 SDN_F3 SDN_L2 SDN_M10 SDN_E1 SDN_F5 SDN_L3 SDN_E6 SDN_E4 SDN_T6 SDN_M8 SDN_L8 SDN_E9
SDN Prefix
SDN_E
SDN_F
SDN_L
SDN_M
SDN_T
Proportion
1
0.5
0
D
Moran's I
0.6
0.4
0.2
0.0
sSDN3 sSDN4 sSDN2 sSDN6 sSDN1 sSDN7 sSDN5
E
Moran's I
0.6
0.4
0.2
0.0
SDN prefix
SDN_T
SDN_E
SDN_F
SDN_L
SDN_M
SDN_M10 SDN_M02 SDN_L2 SDN_T1 SDN_M01 SDN_T6 SDN_L1 SDN_L5 SDN_T3 SDN_T7 SDN_T2 SDN_F3 SDN_T8 SDN_F1 SDN_T5 SDN_M07 SDN_E7 SDN_L3 SDN_M04 SDN_L7 SDN_E1 SDN_T4 SDN_M08 SDN_E3 SDN_F5 SDN_M06 SDN_F4 SDN_M03 SDN_E4 SDN_E6 SDN_L8 SDN_F2 SDN_L6 SDN_E5 SDN_M05 SDN_L4 SDN_E8 SDN_E2 SDN_M09 SDN_E9
F
C3
0.017
Proportion
0.9
0.6
0.3
sSDN2 sSDN4
G
C4
1.8e−10
Proportion
0.9
0.6
0.3
sSDN2 sSDN4
H
LCS-585 T
LCS-576 T
LCS-675 T
LCS-634 T

Figure S4

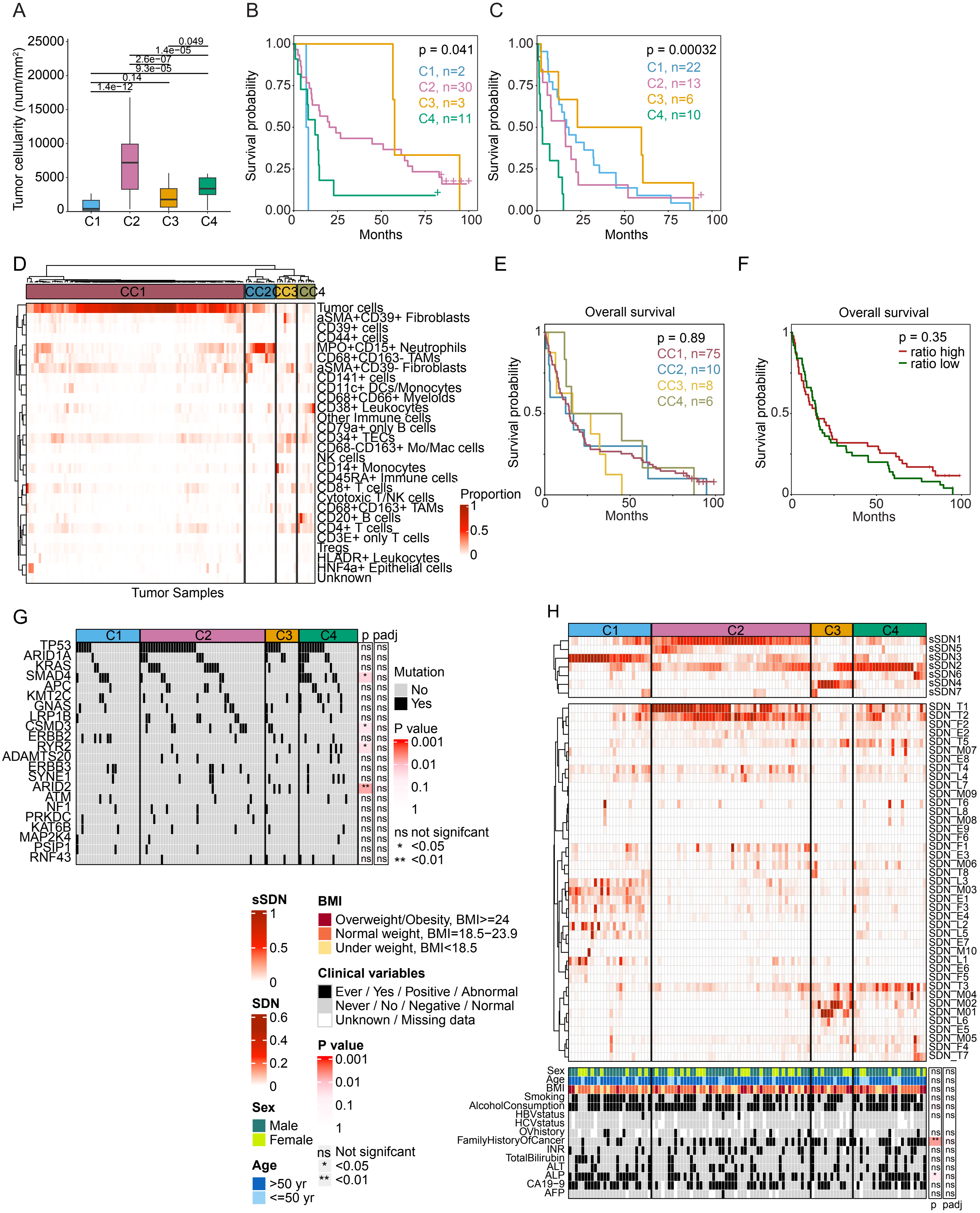

A
B
C
D
E
F
G
H
Tumor cellularity (num/mm²)
C1
C2
C3
C4
p = 0.041
C1, n=2
C2, n=30
C3, n=3
C4, n=11
Survival probability
Months
p = 0.00032
C1, n=22
C2, n=13
C3, n=6
C4, n=10
CC1
CC2
CC3
CC4
Tumor Samples
Proportion
Overall survival
p = 0.89
CC1, n=75
CC2, n=10
CC3, n=8
CC4, n=6
p = 0.35
ratio high
ratio low
Mutation
No
Yes
P value
ns not signifcant
* <0.05
** <0.01
sSDN
SDN
BMI
Overweight/Obesity, BMI>=24
Normal weight, BMI=18.5−23.9
Under weight, BMI<18.5
Clinical variables
Ever / Yes / Positive / Abnormal
Never / No / Negative / Normal
Unknown / Missing data
Sex
Male
Female
Age
>50 yr
<=50 yr
ns Not signifcant

Figure S5

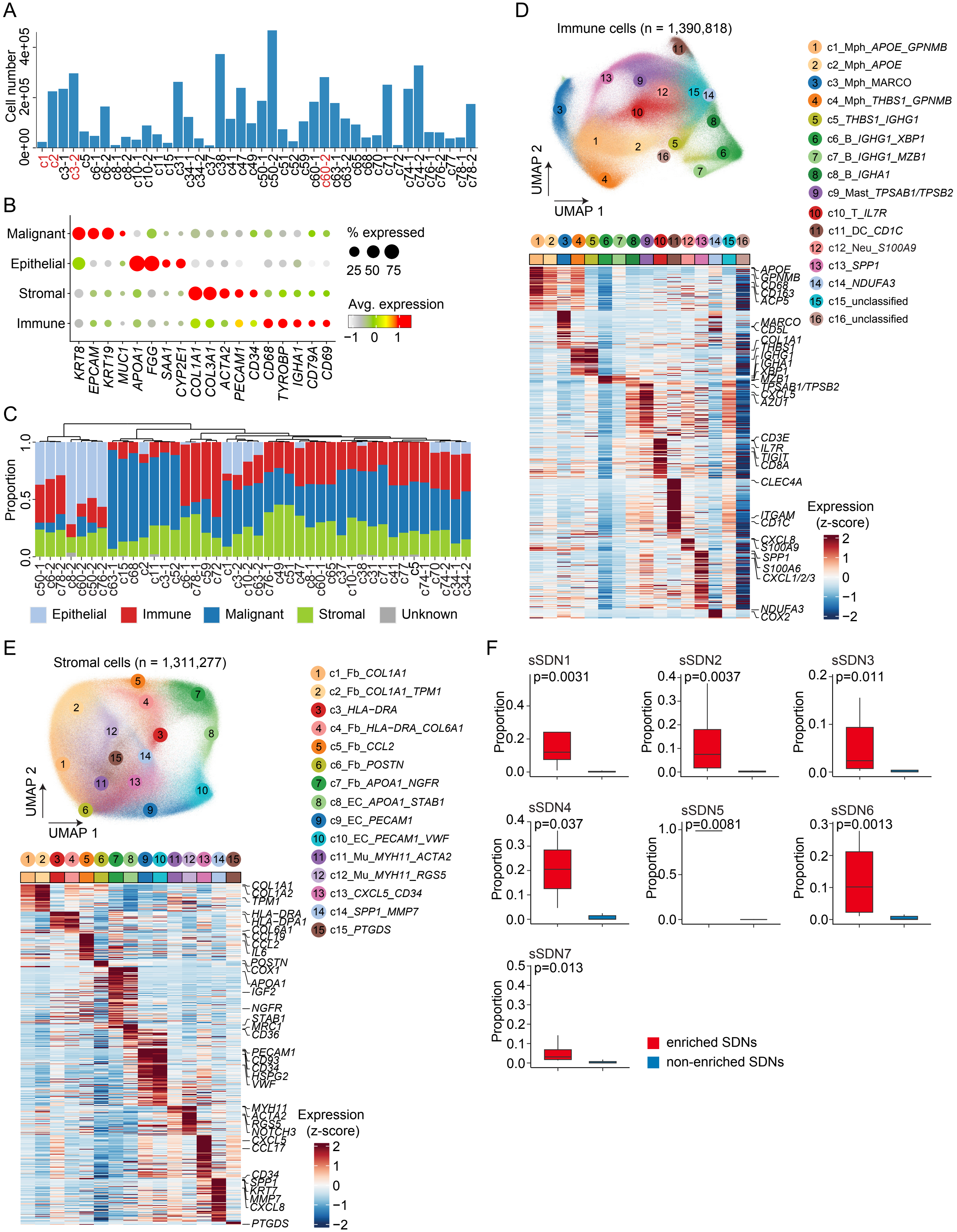

A
Cell number
0e+00
2e+05
4e+05
c1 c2 c3-1 c3-2 c5 c6-1 c6-2 c8-1 c8-2 c10-1 c10-2 c11 c15 c31 c34-1 c34-2 c37 c38 c41 c47 c49 c50-1 c50-2 c51 c52 c59 c60-1 c60-2 c63-1 c63-2 c65 c68 c70 c71 c72 c74-1 c74-2 c76-1 c76-2 c77 c78-1 c78-2
B
Malignant
Epithelial
Stromal
Immune
KRT8 EPCAM KRT19 MUC1 APOA1 FGG SAA1 CYP2E1 COL1A1 COL3A1 ACTA2 PECAM1 CD34 CD68 TYROBP IGHA1 CD79A CD69
% expressed
25 50 75
Avg. expression
−1 0 1
C
Proportion
0.0
0.5
1.0
c50-1 c6-2 c78-2 c8-2 c60-2 c50-2 c76-2 c63-1 c15 c68 c2 c11 c3-1 c52 c6-1 c78-1 c59 c72 c1 c3-2 c10-2 c63-2 c76-1 c49 c51 c47 c8-1 c60-1 c65 c37 c10-1 c38 c31 c71 c41 c77 c5 c74-1 c70 c74-2 c34-1 c34-2
Epithelial
Immune
Malignant
Stromal
Unknown
D
Immune cells (n = 1,390,818)
UMAP 1
UMAP 2
c1_Mph_APOE_GPNMB
c2_Mph_APOE
c3_Mph_MARCO
c4_Mph_THBS1_GPNMB
c5_THBS1_IGHG1
c6_B_IGHG1_XBP1
c7_B_IGHG1_MZB1
c8_B_IGHA1
c9_Mast_TPSAB1/TPSB2
c10_T_IL7R
c11_DC_CD1C
c12_Neu_S100A9
c13_SPP1
c14_NDUFA3
c15_unclassified
c16_unclassified
APOE
GPNMB
CD68
CD163
ACP5
MARCO
CD5L
COL1A1
THBS1
IGHG1
IGHA1
XBP1
MZB1
TPSAB1/TPSB2
CXCL5
AZU1
CD3E
IL7R
TIGIT
CD8A
CLEC4A
ITGAM
CD1C
CXCL8
S100A9
SPP1
S100A6
CXCL1/2/3
NDUFA3
COX2
Expression
(z-score)
2
1
0
−1
−2
E
Stromal cells (n = 1,311,277)
UMAP 1
UMAP 2
c1_Fb_COL1A1
c2_Fb_COL1A1_TPM1
c3_HLA−DRA
c4_Fb_HLA−DRA_COL6A1
c5_Fb_CCL2
c6_Fb_POSTN
c7_Fb_APOA1_NGFR
c8_EC_APOA1_STAB1
c9_EC_PECAM1
c10_EC_PECAM1_VWF
c11_Mu_MYH11_ACTA2
c12_Mu_MYH11_RGS5
c13_CXCL5_CD34
c14_SPP1_MMP7
c15_PTGDS
COL1A1
COL1A2
TPM1
HLA−DRA
HLA−DPA1
COL6A1
CCL19
CCL2
IL6
POSTN
COX1
APOA1
IGF2
NGFR
STAB1
MRC1
CD36
PECAM1
CD93
CD34
HSPG2
VWF
MYH11
ACTA2
RGS5
NOTCH3
CXCL5
CCL17
CD34
SPP1
KRT7
MMP7
CXCL8
PTGDS
Expression
(z-score)
2
1
0
−1
−2
F
sSDN1 p=0.0031
sSDN2 p=0.0037
sSDN3 p=0.011
sSDN4 p=0.037
sSDN5 p=0.0081
sSDN6 p=0.0013
sSDN7 p=0.013
Proportion
enriched SDNs
non-enriched SDNs

Figure S6

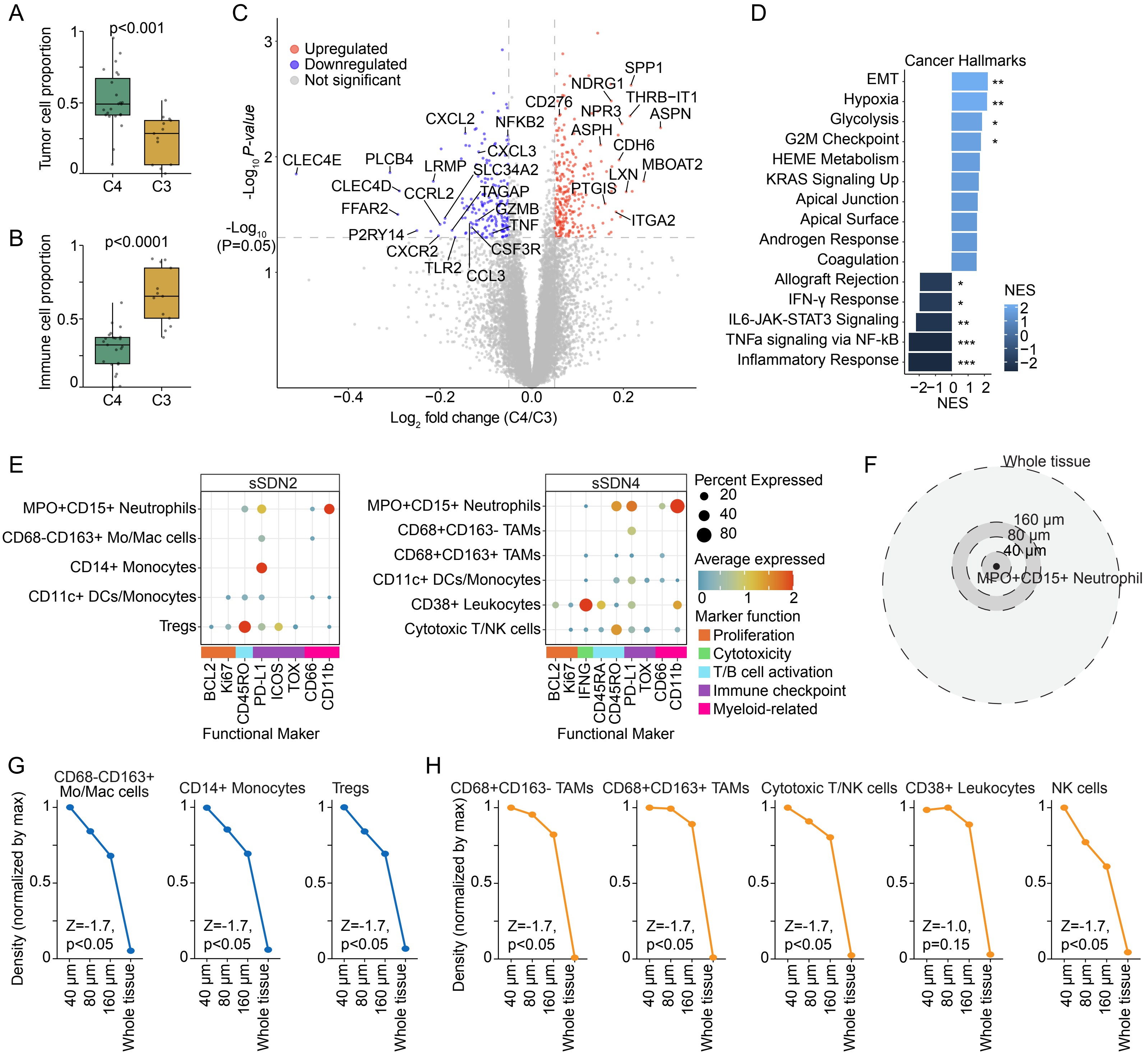
A
p<0.001
Tumor cell proportion
1
0.5
0
C4
C3
B
p<0.0001
Immune cell proportion
1
0.5
0
C4
C3
C
Upregulated
Downregulated
Not significant
-Log10 P-value
-Log10 (P=0.05)
Log2 fold change (C4/C3)
−0.4
−0.2
0.0
0.2
3
2
1
CLEC4E
PLCB4
LRMP
CXCL2
NFKB2
CXCL3
SLC34A2
CLEC4D
CCRL2
FFAR2
TAGAP
GZMB
TNF
P2RY14
CXCR2
CSF3R
TLR2
CCL3
SPP1
NDRG1
THRB−IT1
CD276
NPR3
ASPN
ASPH
CDH6
LXN
MBOAT2
PTGIS
ITGA2
D
Cancer Hallmarks
EMT
Hypoxia
Glycolysis
G2M Checkpoint
HEME Metabolism
KRAS Signaling Up
Apical Junction
Apical Surface
Androgen Response
Coagulation
Allograft Rejection
IFN-γ Response
IL6-JAK-STAT3 Signaling
TNFa signaling via NF-kB
Inflammatory Response
**
**
*
*
*
*
**
***
***
NES
2
1
0
−1
−2
−2−1 0 1 2
NES
E
sSDN2
MPO+CD15+ Neutrophils
CD68-CD163+ Mo/Mac cells
CD14+ Monocytes
CD11c+ DCs/Monocytes
Tregs
BCL2
Ki67
CD45RO
PD-L1
ICOS
TOX
CD66
CD11b
Functional Maker
sSDN4
MPO+CD15+ Neutrophils
CD68+CD163- TAMs
CD68+CD163+ TAMs
CD11c+ DCs/Monocytes
CD38+ Leukocytes
Cytotoxic T/NK cells
BCL2
Ki67
IFNG
CD45RA
CD45RO
PD-L1
TOX
CD66
CD11b
Functional Maker
Percent Expressed
20
40
80
Average expressed
0
1
2
Marker function
Proliferation
Cytotoxicity
T/B cell activation
Immune checkpoint
Myeloid-related
F
Whole tissue
160 μm
80 μm
40 μm
MPO+CD15+ Neutrophil
G
CD68-CD163+ Mo/Mac cells
CD14+ Monocytes
Tregs
Density (normalized by max)
1
0.5
0
Z=-1.7, p<0.05
Z=-1.7, p<0.05
Z=-1.7, p<0.05
40 μm
80 μm
160 μm
Whole tissue
H
CD68+CD163- TAMs
CD68+CD163+ TAMs
Cytotoxic T/NK cells
CD38+ Leukocytes
NK cells
Density (normalized by max)
Z=-1.7, p<0.05
Z=-1.7, p<0.05
Z=-1.7, p<0.05
Z=-1.0, p=0.15
Z=-1.7, p<0.05
40 μm
80 μm
160 μm
Whole tissue

Figure S7

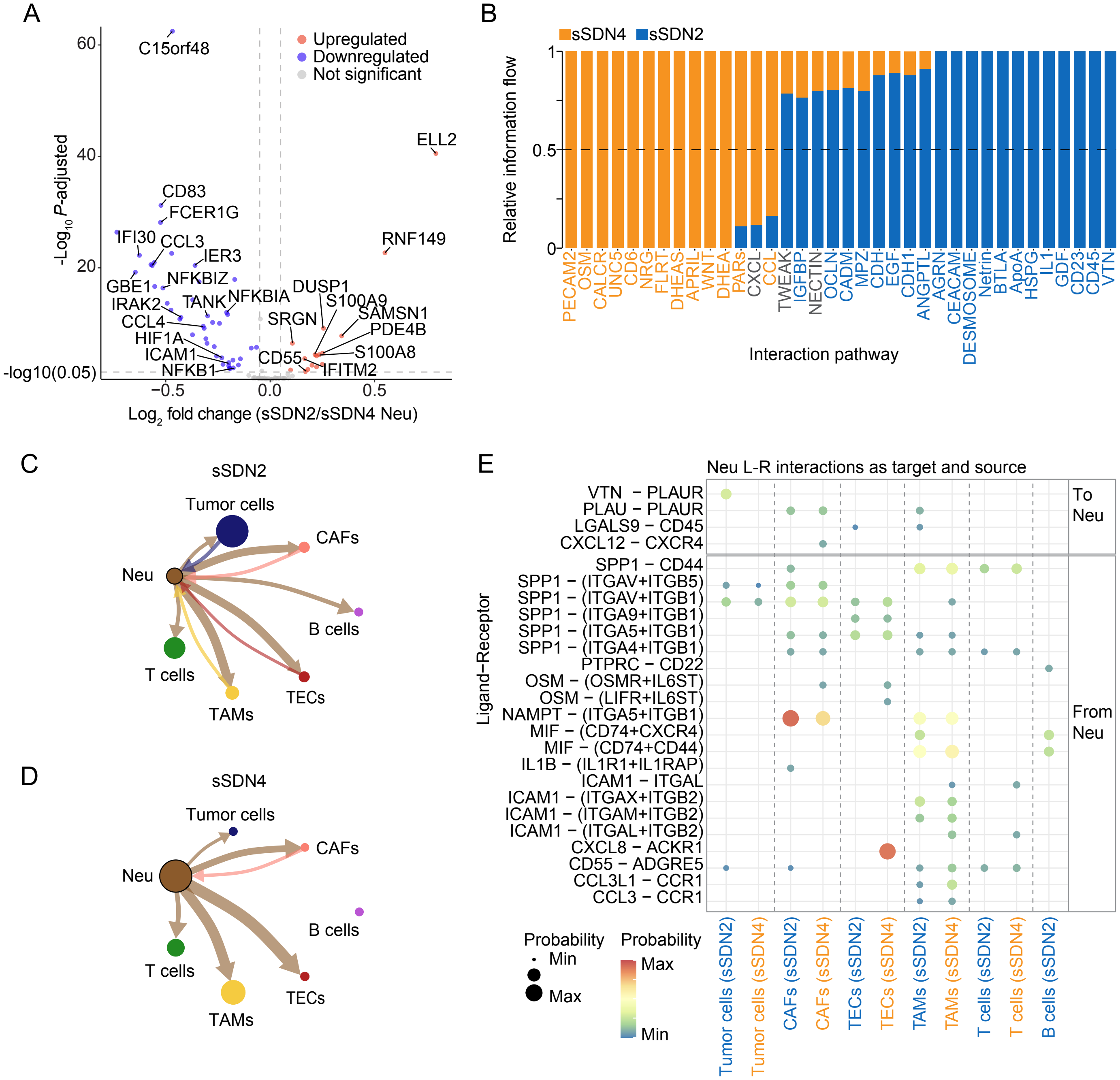
A
Upregulated
Downregulated
Not significant
C15orf48
ELL2
CD83
FCER1G
IFI30
CCL3
IER3
RNF149
GBE1
NFKBIZ
IRAK2
TANK
NFKBIA
DUSP1
S100A9
CCL4
SRGN
SAMSN1
PDE4B
HIF1A
ICAM1
CD55
S100A8
NFKB1
IFITM2
-Log10 P-adjusted
-log10(0.05)
Log2 fold change (sSDN2/sSDN4 Neu)
B
sSDN4
sSDN2
Relative information flow
Interaction pathway
PECAM2
OSM
CALCR
UNC5
CD6
NRG
FLRT
DHEAS
APRIL
WNT
DHEA
PARs
CXCL
CCL
TWEAK
IGFBP
NECTIN
OCLN
CADM
MPZ
CDH
EGF
CDH1
ANGPTL
AGRN
CEACAM
DESMOSOME
Netrin
BTLA
ApoA
HSPG
IL1
GDF
CD23
CD45
VTN
C
sSDN2
Tumor cells
CAFs
Neu
B cells
T cells
TECs
TAMs
D
sSDN4
Tumor cells
CAFs
Neu
B cells
T cells
TECs
TAMs
E
Neu L-R interactions as target and source
Ligand−Receptor
VTN − PLAUR
PLAU − PLAUR
LGALS9 − CD45
CXCL12 − CXCR4
To Neu
SPP1 − CD44
SPP1 − (ITGAV+ITGB5)
SPP1 − (ITGAV+ITGB1)
SPP1 − (ITGA9+ITGB1)
SPP1 − (ITGA5+ITGB1)
SPP1 − (ITGA4+ITGB1)
PTPRC − CD22
OSM − (OSMR+IL6ST)
OSM − (LIFR+IL6ST)
NAMPT − (ITGA5+ITGB1)
MIF − (CD74+CXCR4)
MIF − (CD74+CD44)
IL1B − (IL1R1+IL1RAP)
ICAM1 − ITGAL
ICAM1 − (ITGAX+ITGB2)
ICAM1 − (ITGAM+ITGB2)
ICAM1 − (ITGAL+ITGB2)
CXCL8 − ACKR1
CD55 − ADGRE5
CCL3L1 − CCR1
CCL3 − CCR1
From Neu
Probability
Min
Max
Probability
Max
Min
Tumor cells (sSDN2)
Tumor cells (sSDN4)
CAFs (sSDN2)
CAFs (sSDN4)
TECs (sSDN2)
TECs (sSDN4)
TAMs (sSDN2)
TAMs (sSDN4)
T cells (sSDN2)
T cells (sSDN4)
B cells (sSDN2)

Figure S8

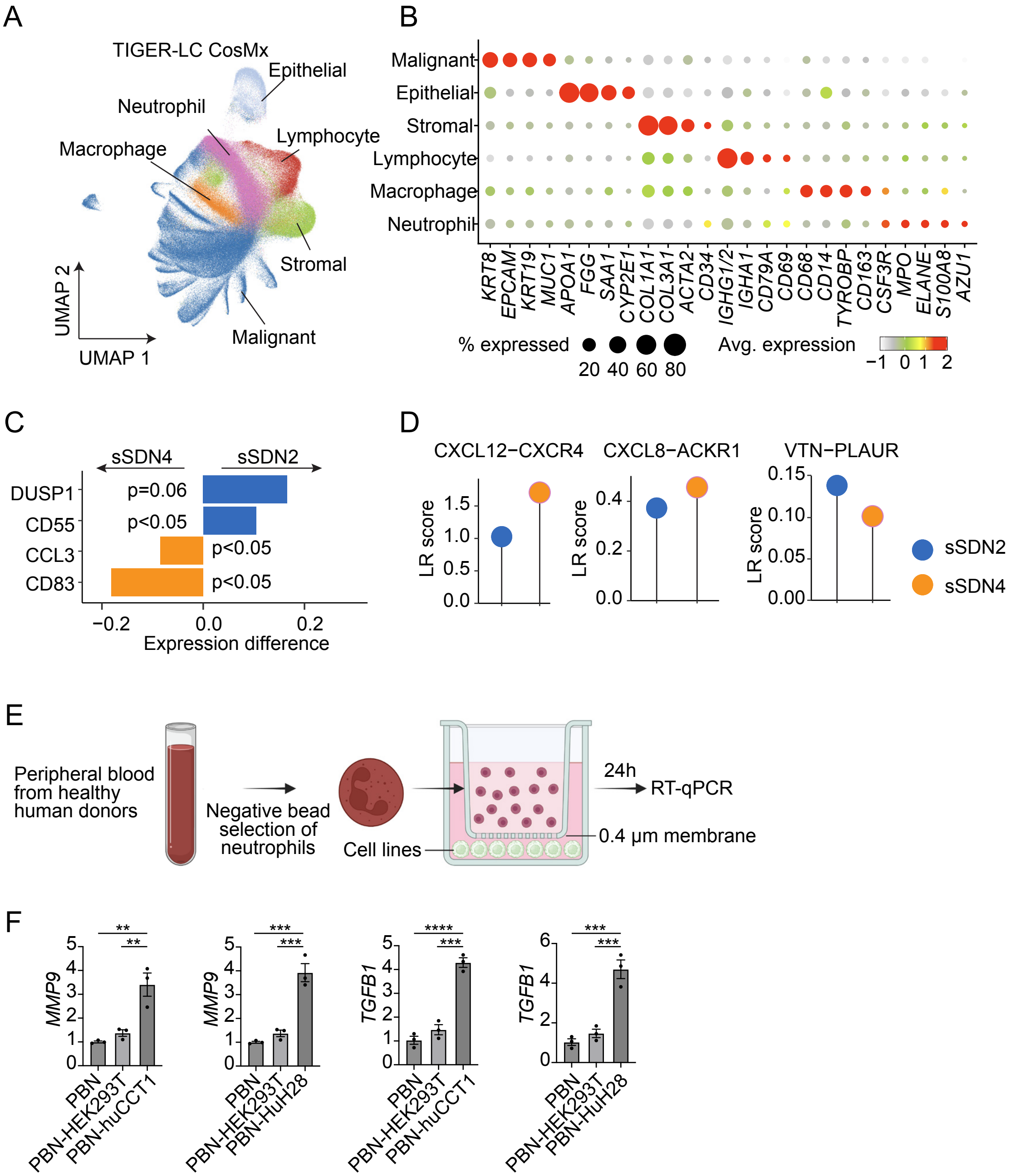

A
TIGER-LC CosMx
Epithelial
Neutrophil
Lymphocyte
Macrophage
Stromal
Malignant
UMAP 2
UMAP 1
B
Malignant
Epithelial
Stromal
Lymphocyte
Macrophage
Neutrophil
KRT8
EPCAM
KRT19
MUC1
APOA1
FGG
SAA1
CYP2E1
COL1A1
COL3A1
ACTA2
CD34
IGHG1/2
IGHA1
CD79A
CD69
CD68
CD14
TYROBP
CD163
CSF3R
MPO
ELANE
S100A8
AZU1
% expressed
20 40 60 80
Avg. expression
−1 0 1 2
C
sSDN4
sSDN2
DUSP1
CD55
CCL3
CD83
p=0.06
p<0.05
p<0.05
p<0.05
−0.2 0.0 0.2
Expression difference
D
CXCL12−CXCR4
CXCL8−ACKR1
VTN−PLAUR
LR score
sSDN2
sSDN4
E
Peripheral blood from healthy human donors
Negative bead selection of neutrophils
Cell lines
24h
RT-qPCR
0.4 μm membrane
F
MMP9
TGFB1
PBN
PBN-HEK293T
PBN-huCCT1
PBN-HuH28

Figure S9

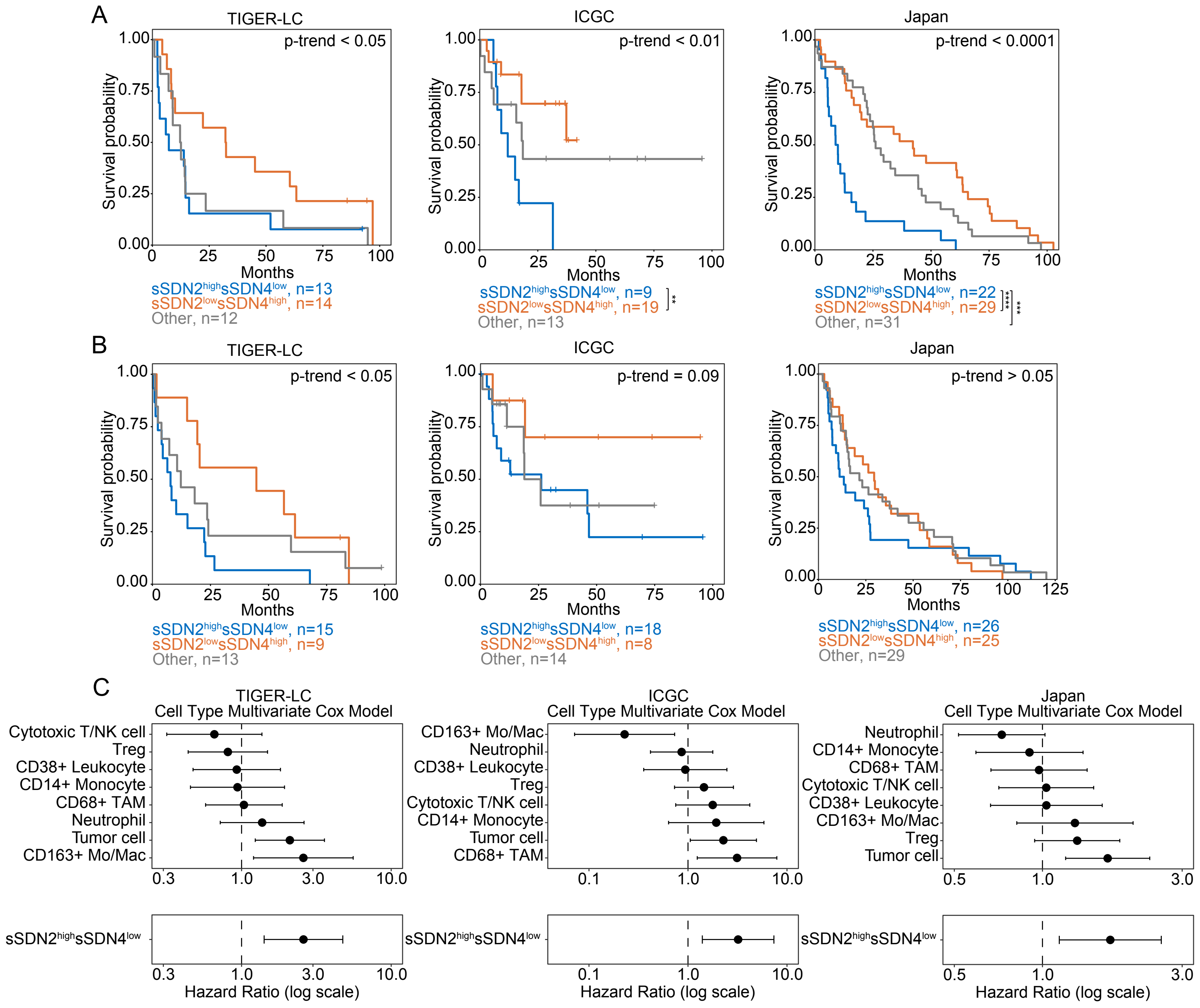
A
TIGER-LC
p-trend < 0.05
ICGC
p-trend < 0.01
Japan
p-trend < 0.0001
Survival probability
Months
sSDN2highsSDN4low, n=13
sSDN2lowsSDN4high, n=14
Other, n=12
sSDN2highsSDN4low, n=9
sSDN2lowsSDN4high, n=19
Other, n=13
sSDN2highsSDN4low, n=22
sSDN2lowsSDN4high, n=29
Other, n=31
B
TIGER-LC
p-trend < 0.05
ICGC
p-trend = 0.09
Japan
p-trend > 0.05
Survival probability
Months
sSDN2highsSDN4low, n=15
sSDN2lowsSDN4high, n=9
Other, n=13
sSDN2highsSDN4low, n=18
sSDN2lowsSDN4high, n=8
Other, n=14
sSDN2highsSDN4low, n=26
sSDN2lowsSDN4high, n=25
Other, n=29
C
TIGER-LC
Cell Type Multivariate Cox Model
Cytotoxic T/NK cell
Treg
CD38+ Leukocyte
CD14+ Monocyte
CD68+ TAM
Neutrophil
Tumor cell
CD163+ Mo/Mac
ICGC
Cell Type Multivariate Cox Model
CD163+ Mo/Mac
Neutrophil
CD38+ Leukocyte
Treg
Cytotoxic T/NK cell
CD14+ Monocyte
Tumor cell
CD68+ TAM
Japan
Cell Type Multivariate Cox Model
Neutrophil
CD14+ Monocyte
CD68+ TAM
Cytotoxic T/NK cell
CD38+ Leukocyte
CD163+ Mo/Mac
Treg
Tumor cell
sSDN2highsSDN4low
Hazard Ratio (log scale)

**Table S1. Demographic, clinical, and laboratory characteristics of iCCA patients in the TIGER-LC cohorts at the time of surgery.**

| Clinical variable | Patients (n = 131) |
|---|---|
| **Demographic–no. (%)** | |
| **Sex** | |
| Male | 88 (67) |
| Female | 43 (33) |
| **Age** | |
| >50 yr | 111 (85) |
| ≤50 yr | 20 (15) |
| **Risk factors–no. (%)** | |
| **BMI** | |
| Normal weight, BMI = 18.5–23.9 | 77 (59) |
| Overweight/Obesity, BMI ≥24 | 28 (21) |
| Under weight, BMI <18.5 | 26 (20) |
| **Smoking** | |
| Never | 49 (37) |
| Ever | 81 (62) |
| Missing data | 1 (1) |
| **Alcohol consumption** | |
| No | 40 (31) |
| Yes | 91 (69) |
| **HBV status** | |
| Positive | 4 (3) |
| Negative | 104 (79) |
| Missing data | 23 (18) |
| **HCV status** | |
| Positive | 0 (0) |
| Negative | 107 (82) |
| Missing data | 24 (18) |
| **History of OV** | |
| No | 84 (64) |
| Yes | 15 (11) |
| Unknown | 30 (23) |
| Missing data | 2 (2) |
| **Family history of cancer** | |
| No | 74 (56) |
| Yes | 49 (37) |
| Unknown | 8 (6) |
| **Liver function factors–no. (%)** | |
| **Child-Pugh class** | |
| A | 64 (49) |
| B | 8 (6) |
| C | 3 (2) |
| Missing data | 56 (43) |
| **INR (International normalized ratio)** | |
| Normal, <1.1 | 90 (69) |
| Abnormal, ≥1.1 | 40 (31) |

| | |
|---|---|
| Missing data | 1 (1) |
| **Total bilirubin** | |
| Normal, ≤1.9mg/dL | 101 (77) |
| Abnormal, >1.9mg/dL | 30 (23) |
| **ALT (Alanine aminotransferase)** | |
| Normal, ≤50U/L | 93 (71) |
| Abnormal, >50U/L | 38 (29) |
| **ALP (Alkaline phosphatase)** | |
| Normal, ≤147U/L | 58 (44) |
| Abnormal, >147U/L | 73 (56) |
| **CA19-9** | |
| Normal, ≤37U/mL | 48 (37) |
| Abnormal, >37U/mL | 76 (58) |
| Missing data | 7 (5) |
| **AFP (Alpha-fetoprotein)** | |
| Normal, ≤300 ng/mL | 110 (84) |
| Abnormal, >300 ng/mL | 2 (2) |
| Missing data | 19 (15) |
| **Tumor features–no. (%)** | |
| **Tumor size** | |
| ≤5 cm | 45 (34) |
| >5 cm | 45 (34) |
| Missing data | 41 (31) |
| **Multinodular tumor** | |
| No | 63 (48) |
| Yes | 21 (16) |
| Missing data | 47 (36) |
| **TNM staging** | |
| I | 16 (12) |
| II | 8 (6) |
| III-IV | 46 (35) |
| Missing data | 61 (47) |
| **Distal metastasis** | |
| No | 44 (34) |
| Yes | 11 (8) |
| Missing data | 76 (58) |
| **Perineural invasion** | |
| Yes | 24 (18) |
| No | 26 (20) |
| Missing data | 81 (62) |
| **Lymphatic invasion** | |
| Yes | 31 (24) |
| No | 19 (15) |
| Missing data | 81 (62) |
| **Portaltract invasion** | |
| Yes | 27 (21) |
| No | 23 (18) |
| Missing data | 81 (62) |
| **Ductal dysplasia** | |

| | |
|---|---|
| Yes | 24 (18) |
| No | 26 (20) |
| Missing data | 81 (62) |
| **Stromal infiltration** | |
| Yes | 32 (24) |
| No | 18 (14) |
| Missing data | 81 (62) |

**Table S2. Antibodies used for CODEX staining of iCCA in this study.**

| Antibody | Source | Clone | Vendor | Catalogue number | Dilution |
|---|---|---|---|---|---|
| aSMA | custom | 1A4 | BioRad | MCA5781GA | 1:120 |
| BCL2 | commercial | EPR17509 | Akoya Biosciences | 4250098 | 1:200 |
| b-catenin | commercial | 12F7 | Akoya Biosciences | 4250091 | 1:200 |
| Caveolin1 | commercial | D46G3 | Akoya Biosciences | 4450084 | 1:200 |
| CD107a | commercial | H4A3 | Akoya Biosciences | 4550098 | 1:200 |
| CD11b | custom | EPR1344 | Abcam | ab209970 | 1:80 |
| CD11c | custom | EP1347Y | Abcam | ab216655 | 1:80 |
| CD14 | commercial | EPR3653 | Akoya Biosciences | 4450047 | 1:200 |
| CD141 | commercial | E7Y9P | Akoya Biosciences | 4250097 | 1:200 |
| CD15 | custom | HI-98 | Biolegend | 301902 | 1:150 |
| CD163 | commercial | EPR19518 | Akoya Biosciences | 4250079 | 1:200 |
| CD20 | commercial | L26 | Akoya Biosciences | 4150018 | 1:200 |
| CD21 | commercial | EP3093 | Akoya Biosciences | 4450100 | 1:200 |
| CD3E | commercial | EP449E | Akoya Biosciences | 4450030 | 1:200 |
| CD31 | custom | EP3095 | Abcam | ab226157 | 1:100 |
| CD34 | custom | QBEnd/10 | BioTechne | NBP2-34713 | 1:100 |
| CD38 | commercial | E7Z8C | Akoya Biosciences | 4250080 | 1:200 |
| CD39 | commercial | EPR20627 | Akoya Biosciences | 4250076 | 1:200 |
| CD4 | commercial | EPR6855 | Akoya Biosciences | 4350018 | 1:200 |
| CD44 | commercial | 156-3C11 | Akoya Biosciences | 4450041 | 1:200 |
| CD45 | commercial | D9M81 | Akoya Biosciences | 4450042 | 1:200 |
| CD45RA | custom | HI100 | Biolegend | 304102 | 1:100 |
| CD45RO | custom | UCHL1 | Akoya Biosciences | 4250023 | 1:200 |
| CD56 | commercial | CAL53 | Akoya Biosciences | 4250087 | 1:200 |
| CD57 | custom | HNK1 | Biolegend | 359602 | 1:120 |
| CD66 | commercial | ASL-32 | Akoya Biosciences | 4550001 | 1:200 |
| CD68 | commercial | KP1 | Akoya Biosciences | 4350019 | 1:200 |
| CD79a | commercial | D1X5C | Akoya Biosciences | 4450078 | 1:200 |
| CD8 | commercial | C8/144B | Akoya Biosciences | 4250012 | 1:200 |
| E-cadherin | commercial | 4A2C7 | Akoya Biosciences | 4250021 | 1:200 |
| EpCAM | commercial | D9S3P | Akoya Biosciences | 4450088 | 1:200 |
| FOXP3 | commercial | SP97 | Akoya Biosciences | 4550070 | 1:200 |
| GZMB | commercial | D6E9W | Akoya Biosciences | 4250055 | 1:200 |
| HIF1A | commercial | VISTA | Akoya Biosciences | 4550069 | 1:200 |
| HLA-A | commercial | EP1395Y | Akoya Biosciences | 4450046 | 1:200 |
| HLA-DR | commercial | EPR3692 | Akoya Biosciences | 4450029 | 1:200 |
| HLA-E | commercial | EPR25300-104 | Akoya Biosciences | 4250065 | 1:200 |
| HNF4a | custom | K9218 | Thermo Fisher Scientific | MA1-199 | 1:100 |
| ICOS | commercial | D1K2T | Akoya Biosciences | 4350059 | 1:200 |
| IFNG | commercial | EPR21704 | Akoya Biosciences | 4250062 | 1:200 |
| Ki67 | commercial | B56 | Akoya Biosciences | 4250019 | 1:200 |
| LAG3 | commercial | EPR20261 | Akoya Biosciences | 4550058 | 1:100 |

| | | | | | |
|---|---|---|---|---|---|
| MPO | commercial | E1E7I | Akoya Biosciences | 4250083 | 1:200 |
| NaKATPase | custom | EP1845Y | Abcam | ab167390 | 1:100 |
| PanCK | commercial | AE-1/AE-3 | Akoya Biosciences | 4450020 | 1:200 |
| PCNA | commercial | PC10 | Akoya Biosciences | 4550124 | 1:200 |
| PD-1 | commercial | D4W2J | Akoya Biosciences | 4550038 | 1:200 |
| PD-L1 | commercial | 73-10 | Akoya Biosciences | 4550072 | 1:200 |
| TOX | commercial | E6I3Q | Akoya Biosciences | 4250067 | 1:200 |
| TP63 | commercial | W15093A | Akoya Biosciences | 4550081 | 1:200 |
| VISTA | commercial | D1L2G | Akoya Biosciences | 4250063 | 1:120 |

**Table S3. CODEX antibody marker panel.**

| CODEX (FF) panel with 51 markers + 2 nuclear stain | | |
|---|---|---|
| **Immune**<br>CD45<br>CD45RA CD45RO CD38<br>**T cells**<br>CD3E CD4 CD8<br>FOXP3<br>**B cells**<br>CD20 CD21 CD79a<br>**NK cells**<br>CD56 CD57<br>**Myeloid cells**<br>CD14 CD11b CD11c<br>CD15 MPO CD66<br>CD68 CD163 | **Epithelial**<br>b-catenin E-cadherin<br>**Hepatocytes**<br>HNF4a<br>**Biliary epithelial**<br>EpCAM PanCK | **Functional**<br>Ki67 PCNA<br>HIF1A<br>BCL2 CD44<br>CD39 CD107a<br>HLA-DR HLA-E*<br>HLA-A$^{+}$<br>IFNG GZMB<br>PD-1 PD-L1<br>ICOS VISTA<br>LAG3 TOX |
| | **Stromal**<br>aSMA* CD31 CD34 | **Nuclear Stain**<br>DAPI DRAQ5 |
| | **Other**<br>CD141<br>Caveolin1<br>NaKATPase<br>TP63* | |

* Only used for tumor samples. $^{+}$Only used for non-tumor samples.

**Table S4. Clinical information of iCCA patients in the European cohort at the time of surgery.**

| Clinical variable | Patients (n = 31) |
|---|---|
| **Demographic–no. (%)** | |
| **Sex** | |
| Male | 13 (42%) |
| Female | 18 (58%) |
| **Age** | |
| ≤50 yr | 5 (16%) |
| >50 yr | 19 (61%) |
| Missing data | 7 (23%) |
| **Risk factors–no. (%)** | |
| **HBV status** | |
| Positive | 0 (0%) |
| Negative | 29 (94%) |
| Missing data | 2 (6.5%) |
| **HCV status** | |
| Positive | 0 (0%) |
| Negative | 29 (94%) |
| Missing data | 2 (6.5%) |
| **Liver function factors–no. (%)** | |
| **Total bilirubin** | |
| Normal, ≤1.9 mg/dL | 21 (68%) |
| Abnormal, >1.9 mg/dL | 3 (9.7%) |
| Missing data | 7 (23%) |
| **ALT (Alanine aminotrasferase)** | |
| Normal, ≤50 U/L | 14 (45%) |
| Abnormal, >50 U/L | 7 (23%) |
| Missing data | 10 (32%) |
| **ALP (Alkaline phosphatase)** | |
| Normal, ≤147 U/L | 11 (35%) |
| Abnormal, >147 U/L | 6 (19%) |
| Missing data | 14 (45%) |
| **CA19-9** | |
| Normal, ≤37 U/mL | 8 (26%) |
| Abnormal, >37 U/mL | 7 (23%) |
| Missing data | 16 (52%) |
| **Tumor features–no. (%)** | |
| **Tumor size** | |
| ≤5 cm | 8 (26%) |
| >5 cm | 23 (74%) |
| **Multinodular tumor** | |
| Yes | 0 (0%) |
| No | 31 (100%) |
| **TNM staging** | |
| I | 11 (35%) |

| | |
|---|---|
| II | 9 (29%) |
| III | 8 (26%) |
| Missing data | 3 (9.7%) |
| **Distal metastasis** | |
| Yes | 0 (0%) |
| No | 28 (90%) |
| Missing data | 3 (9.7%) |
| **Vascular invasion** | |
| Yes | 2 (6.5%) |
| No | 20 (65%) |
| Missing data | 9 (29%) |
